# Beyond range size: drivers of species’ geographic range structure in European plants

**DOI:** 10.1101/2020.02.08.939819

**Authors:** Anna M. Csergő, Olivier Broennimann, Antoine Guisan, Yvonne M. Buckley

**Affiliations:** School of Natural Sciences, Zoology, Trinity College Dublin, The University of Dublin, Dublin 2, Ireland; Department of Botany and Soroksár Botanical Garden, Szent István University, 1118 Budapest, Hungary; Department of Ecology and Evolution, University of Lausanne, 1015 Lausanne, Switzerland; Institute of Earth Surface Dynamics, University of Lausanne, 1015 Lausanne, Switzerland; ARC Centre of Excellence for Environmental Decisions, School of Biological Sciences, The University of Queensland, Queensland 4072, Australia

**Keywords:** Connectivity, Ecological Niche Model, Europe, Endemic plant species, Fragmentation, Geographic range size, Habitat suitability, Species’ area, Species Distribution Model, Landscape Metrics

## Abstract

**Aim:** To assess if and how species’ range size relates to range structure, if the observed geographic range properties can be retrieved from predicted maps based on species distribution modeling, and whether range properties are predictable from biogeophysical factors.

**Location:** Europe

**Time period:** Current

**Major taxa studied:** 813 vascular plant species endemic to Europe

**Methods:** We quantified the size and spatial structure of species’ geographic ranges and compared ranges currently occupied with those predicted by species distribution models (SDMs). SDMs were constructed using complete occurrence data from the Atlas Florae Europaeae and climatic, soil and topographic predictors. We used landscape metrics to characterize range size, range division and patch shape structure, and analysed the phylogenetic, geographic and ecological drivers of species’ range size and structure using phylogenetic generalized least squares (pGLS).

**Results:** Range structure metrics were mostly decoupled from species’ range size. We found large differences in range metrics between observed and predicted ranges, in particular for species with intermediate observed range size and occupied area, and species with low and high observed patch size distribution, geographic range filling, patch shape complexity and geographic range fractality. Elevation heterogeneity, proximity to continental coasts, Southerly or Easterly geographic range positions and narrow ecological niche breadth constrained species’ observed range size and range structure to different extents. The strength and direction of the relationships differed between observed and predicted ranges.

**Main conclusions:** Several range structure metrics, in addition to range size, are needed to adequately describe and understand species’ ranges. Species’ range structure can be well explained by geophysical factors and species niche width, albeit not consistently for observed and predicted ranges. As range structure can have important ecological and evolutionary consequences, we highlight the need to develop better predictive models of range structure than provided by current SDMs, and we identify the kinds of species for which this is most necessary.

## Introduction

Understanding what drives observed large-scale distribution patterns is key to predicting species’ future geographic ranges and vulnerability to global change (Broennimann et al. 2006). Analyses of species’ geographic ranges have mostly focused on determinants of range size (Morueta-Holme et al. 2013, Sheth et al. 2020). However, examining occupancy patterns throughout a species’ range may enable a much better understanding of dispersal and extinction dynamics in response to global environmental changes (Crooks et al. 2011, Cianfrani et al. 2018). While occupancy patterns have been extensively used in landscape-level distribution analyses (Wintle et al. 2018), they have only recently and incompletely been applied to species’ whole ranges (Brown et al. 1996, Crooks et al. 2011, Pearson et al. 2014, Pagel et al. 2019).

Complex interactions and feedbacks between geophysical, ecological and evolutionary processes determine how occupied areas within species’ ranges are structured and connected (Gaston 2003, Martinez-Meyer et al. 2012). A species’ internal range structure is a consequence of the availability and the spatial arrangement of sites that fulfil the requirements of a species’ fundamental niche (Hutchinson’s duality; Colwell and Rangel 2009) together with dispersal and persistence dynamics and biotic interactions (Soberon 2007; Guisan et al. 2017, Pagel et al. 2019). In turn, constraints due to range structure can modulate a whole suite of ecological and evolutionary processes, such as the routes of continental-scale migrations driven by climatic changes (Hampe and Jump 2011), (meta-)population colonization-extinction dynamics (Manna 2017), variation in abundance and genetic diversity within the range (Holt et al. 2002, Pironon et al. 2016), island effects (Covas 2012), or the Allee effect (Lamont et al. 2003). Ultimately, species’ internal range structure can foster divergence among populations and provide opportunities for local adaptation, which might in turn alter species’ fundamental niche and its spatial expression (Colwell & Rangel 2009). Despite the importance of range structure for ecological and evolutionary processes, quantitative analyses of coarse-scale range changes still largely ignore, or insufficiently account for, this property of species’ ranges (but see Crooks et al 2011, Pearson et al. 2014, Cianfrani et al. 2018).

Quantifying geographic range structure requires unbiased knowledge of species distributions (i.e. consistent throughout the range), which is currently unavailable for most organisms. However, the growing biodiversity repositories and species distribution modeling techniques (SDMs, also called ecological niche or habitat suitability models; Guisan et al. 2017) can provide geographically consistent predictions of species’ range-wide distributions. In addition, SDMs can be used to predict not only current but also future range structures, enabling us to move beyond classical range size analyses of climate change impacts (Williams et al. 2008, Cianfrani et al. 2018). A limitation of the approach, however, is that the occupancy of potentially suitable ranges is often incomplete for some species, depending on range shift histories, human land use, dispersal limitations, metapopulation dynamics and species’ other biological properties (Svenning et al. 2006, Estrada et al. 2015, Miller and McGill 2017), leading to mismatches between observed and SDM-predicted geographic patterns. Surprisingly, the ability of SDMs to reproduce species’ observed internal range geometry has not been thoroughly tested to date. One reason for such limitation might come from the lack of comprehensive distribution data at a large scale to develop models and test predictions. The Atlas Florae Europaeae (AFE) is one of the few datasets with coarse-grained, yet reasonably non-biased presence-absence data across the entire range of a large number of plant species in Europe. Here, we developed SDMs for over 800 endemic vascular plant species in AFE and used observed and predicted distributions to quantify species’ geographic range size and structure and explore their potential intrinsic and extrinsic determinants. Specifically, across all species, we addressed the following questions: 1) Is range size coupled with range structure? 2) Are measures of range size and structure congruent between observed and predicted ranges? 3) What niche - related and extrinsic biogeophysical factors explain range size and range structure, and are they the same for measures based on observations and predictions?

## Methods

### Species selection

We accessed species’ occurrence data at 50×50 km^2^ resolution from the database of Atlas Florae Europaeae (AFE) which contains 4 473 taxa from 78 vascular plant families. We subset the database to include European endemic species using EvaplantE, the Database on Endemic Vascular Plants in Europe (Hobohm 2014), which we complemented with our own search for species considered endemic by Flora Europaea using the online search engine of Flora Europaea, with the visual aid of biogeographic atlases (Meusel and Jäger, 1992, digitized: http://chorologie.biologie.uni-halle.de//choro/). This preselection resulted in 1401 taxa. To decrease the risk of bias in SDM predictions from low number of occurrences (Wisz et al. 2008), we ran a second selection to include only taxa with at least 10 occurrence data points, resulting in 864 taxa. We removed 14 species which were unavailable in the phylogenetic tree, and further 37 species because the shape of the minimum convex polygon impeded the calculation of particular range metrics, resulting in a final dataset of 813 species. The taxonomy and nomenclature follows the database of the AFE.

### Species distribution models (range modelling)

To characterise environmental and geographic variation throughout species’ ranges, we selected eight climate variables from the CliMond Archive (Kriticos et al. 2014), five soil variables from the International Geosphere-Biosphere Programme IGBP (Global Soil Data Task 2014), and one topographic variable calculated from elevation maps (Jarvis et al. 2008), totaling 14 final predictors: mean diurnal temperature range, temperature annual range, mean temperature of the wettest quarter, mean temperature of driest quarter, mean temperature of warmest quarter, annual precipitation, precipitation seasonality, precipitation of driest quarter, nitrogen density (minimum), nitrogen density (maximum), thermal capacity, wilting point (minimum), wilting point (maximum), slope (selection details in Appendix S1). We built Ensemble of Small Models (ESM; Breiner et al. 2015) and classical species distribution models (SDM; Thuiller et al. 2009) to predict habitat suitability for each species using the selected environmental variables (Appendix S1, Dryad Digital Repository).

### Range size and range structure metrics

We calculated two range size metrics: geographic range size and occupied area, and four geographic range structure metrics commonly used in landscape ecology, which we adapted to describe how divided and how irregular geographic ranges are. Range structure metrics included two range division metrics (patch size distribution, geographic range filling) and two patch shape metrics (patch shape complexity, geographic range fractality (Box1, Fig. 1, Appendix S2). These metrics are implemented in FRAGSTATS for ArgGIS (McGarigal et al. 2012) and the *SDMTools* package for R (VanDerWal et al. 2014). We calculated all metrics from maps of observed occurrence and habitat suitability separately, with slight modifications of the original functions (Appendix S2.1). We analysed correlations between different metrics to test for their ability to offer complementary information (Appendix S2.2). We tested the sensitivity of the geographic range size and structure metrics to methodological choices taken to define observed and predicted distributions: **1)** map resolution (raster grain size), **2)** rasterization approaches of the AFE atlas grid, **3)** definition of species’ geographic range boundaries, **4)** species distribution modeling technique, **5)** prediction error of species distribution models, **6)** habitat quality thresholds used to define a species’ potential distribution (Appendices S2.3 to S2.8).

#### Box 1. Geographic range size and range structure terminology

As with the geographic range size measures (Gaston 1996 TREE, Box 1), no standard methodology exists for measuring the structure of species’ geographic ranges. We used the following interpretations:

**Geographic Range Size:** the area of occupied and non-occupied areas within a species’ range boundaries (extent of occurrence, Gaston 1996).

**Occupied Area:** the area of occupied habitat patches (area of occupancy, Gaston 1996)

**Patch Size Distribution:** the subdivision of a species’ geographic range into habitat patches of different sizes relative to species’ geographic range size. Likely reflects processes driving abundance-occupancy patterns (McGarigal et al. 2012, modified).

**Geographic Range Filling:** proportion of a species’ geographic range represented by occupied habitat patches within a species’ geographic range boundaries. Some earlier interpretations of range filling may refer to occupancy of suitable habitats, for which we suggest “suitable habitat occupancy” as a more suggestive terminology. The term range filling proposed here captures differences between species’ geographic range size and occupied area (McGarigal et al. 2012, modified).

**Patch Shape Complexity:** the shape complexity of habitat patches averaged across all habitat patches within a species’ geographic range (McGarigal et al. 2012, modified).

**Geographic Range Fractality:** the consistency of the shape of habitat patches across different patch sizes. Range fractality applied to range-level analyses can be viewed as a general patch shape complexity metric, however in our interpretation it indicates a non-additive range shape complexity i.e., quantification of pattern repetition over different spatial scales (McGarigal et al. 2012, modified).

**Figure 1.**
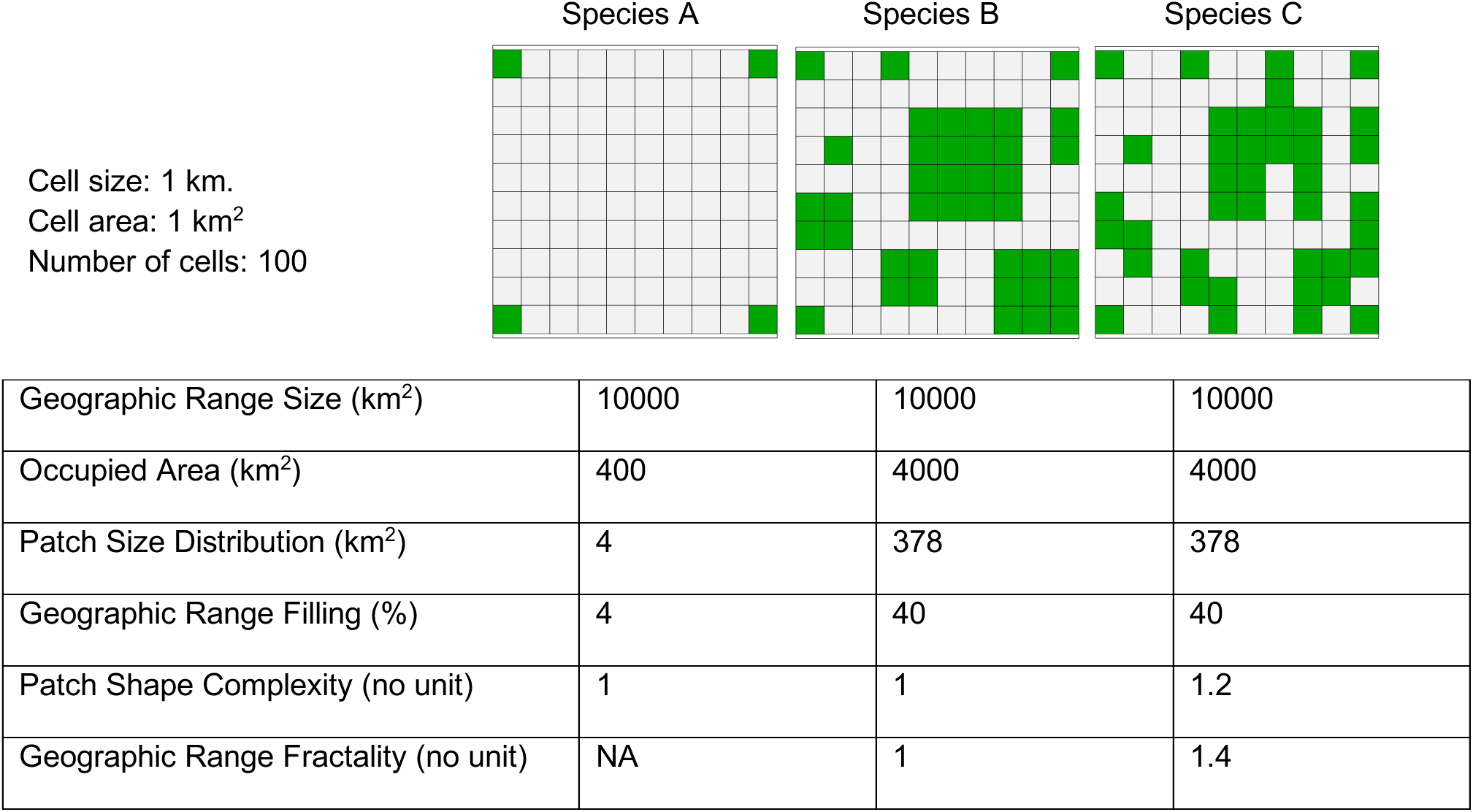
Examples of geographic range size and structure metrics in three theoretical species. The three species have similar geographic range sizes of ca. 10000 km^2^ but differ in their occupied area and range structure metrics. Species A has low occupied area and fills a small proportion of its range, it also has the lowest patch size distribution. The patch shape complexity for A is low. Species B and C have a high and equal occupied area and fill the range to the same extent, with the same patch size distribution. Species B and C differ in their patch shape complexity and range fractality, with C having on average more complex patch shapes, and the complexity of patches increases with patch size. The geographic range fractality cannot be defined for species A, which has a low number of patches.

### Geographic and ecological drivers of range size and structure

We derived geophysical variables and niche breadth that summarise the geographic and ecological properties of species’ ranges. We defined the position of species’ geographic range in Europe as the median latitude and longitude of all 50×50 km2 grids cells occupied (or predicted suitable), and species’ range-wide elevational heterogeneity as the standard deviation of the elevation above sea level across occupied (or predicted suitable) grid cells. We calculated range distance to sea coastline as average distance of each occupied (or predicted suitable) grid cell to the closest shore in Europe, using geographic coordinates in the WGS84 projection map to obtain standardized distances. To calculate niche breadth, we reduced the dimensionality of the climate space (the same climate variables as in the SDM) for all grid cells in Europe to two axes using a PCA analysis (Appendix S3), and we summed the standard deviation of values corresponding to species’ occurrences on each of the axes, weighted by the corresponding axis Eigenvalue.

### Statistical analyses

We first analyzed the relationship between the observed range metrics using Pearson’s r and Spearman’s Rho correlations. Then, to compare species’ range size and structure quantified from observed occurrences and predicted habitat suitability maps respectively, we used phylogenetically uncorrected and corrected paired t-tests (Revell, 2012), using the tree available in Zanne et al. (2014). If any difference between observed and predicted range size and structure metrics was removed upon the phylogenetic correction, that indicated influence of evolutionary history on these differences.

To further test whether the magnitude and direction of differences between observed and predicted suitable range metrics were predictable from the observed range metrics or phylogenetically structured, we fitted Phylogenetic Generalized Least Squares models (pGLS; Freckleton et al. 2002) of the log range metric (*m*) response ratio for predicted (*p*) and occupied (*o*) ranges, with the range metric calculated from observed occurrences (*m_o_*) as the explanatory variable as main and quadratic effects: 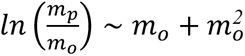. If observed and predicted metrics provide the same information, the response should be constant and set to zero (i.e. ln(1)). By extension, these models indicate for what kinds of observed ranges inferring observed from predicted ranges is the most problematic.

Finally, to test how geographic position, elevation heterogeneity and proximity to seacoast of species’ ranges, as well as species’ niche breadth affected the different range properties, we fitted pGLS models to each of the six observed and predicted metrics separately. The pGLS models were fitted to include the interactions between niche breadth and mean range latitude and between niche breadth and mean range longitude, and the main effects of elevational heterogeneity and proximity to seacoast: *m* ~ *niche breadth* × *latitude* + *niche breadth* × *longitude* + *SD elevation* + *distance to seacoast*. To make effect sizes comparable across different metrics, all explanatory variables were centered on zero with unit variance. To meet model assumptions, geographic range size, occupied (or suitable) area and patch size distribution were log-transformed in all analyses. The phylogenetic t-tests were fitted with the ‘phytools’ package (Revell 2012) and the pGLS models with the ‘caper’ package (Orme et al. 2013) in R 3.2.3 (R Core Development Team 2015).

## Results

### Properties and correlations of observed range metrics

Geographic range metrics quantified from observed distributions were highly variable between species (Fig. 2). As expected, the range size metrics, i.e. geographic range size and occupied area, were highly positively correlated (Pearson’s r = 0.91). Patch size distribution was positively correlated with occupied area (Pearson’s r = 0.682) but only weakly positively correlated with geographic range size (Pearson’s r = 0.34), i.e. large occupied patches added up to larger occupied areas but not necessarily to larger range sizes, and large range sizes were not necessarily composed of large patches. Geographic range filling was negatively correlated with geographic range size (Spearman’s Rho = −0.627), indicating that, in contrast to species with small geographic ranges, species with large geographic ranges had extended unoccupied areas within the species’ range boundaries. Correlations between range structure metrics were generally weak (Pearson’s r or Spearman’s Rho < 0.351), highlighting their complementarity in describing different facets of range structure. An exception was patch size distribution, which was more strongly, positively correlated with geographic range filling, suggesting that species establishing large patches are more likely to fill out species’ ranges. Patch size distribution was also positively correlated with patch shape complexity, indicating spatial constraints on the geometrical complexity of small sized patches (Spearman’s Rho > 0.512) (Appendix S2.2).

**Figure 2.**
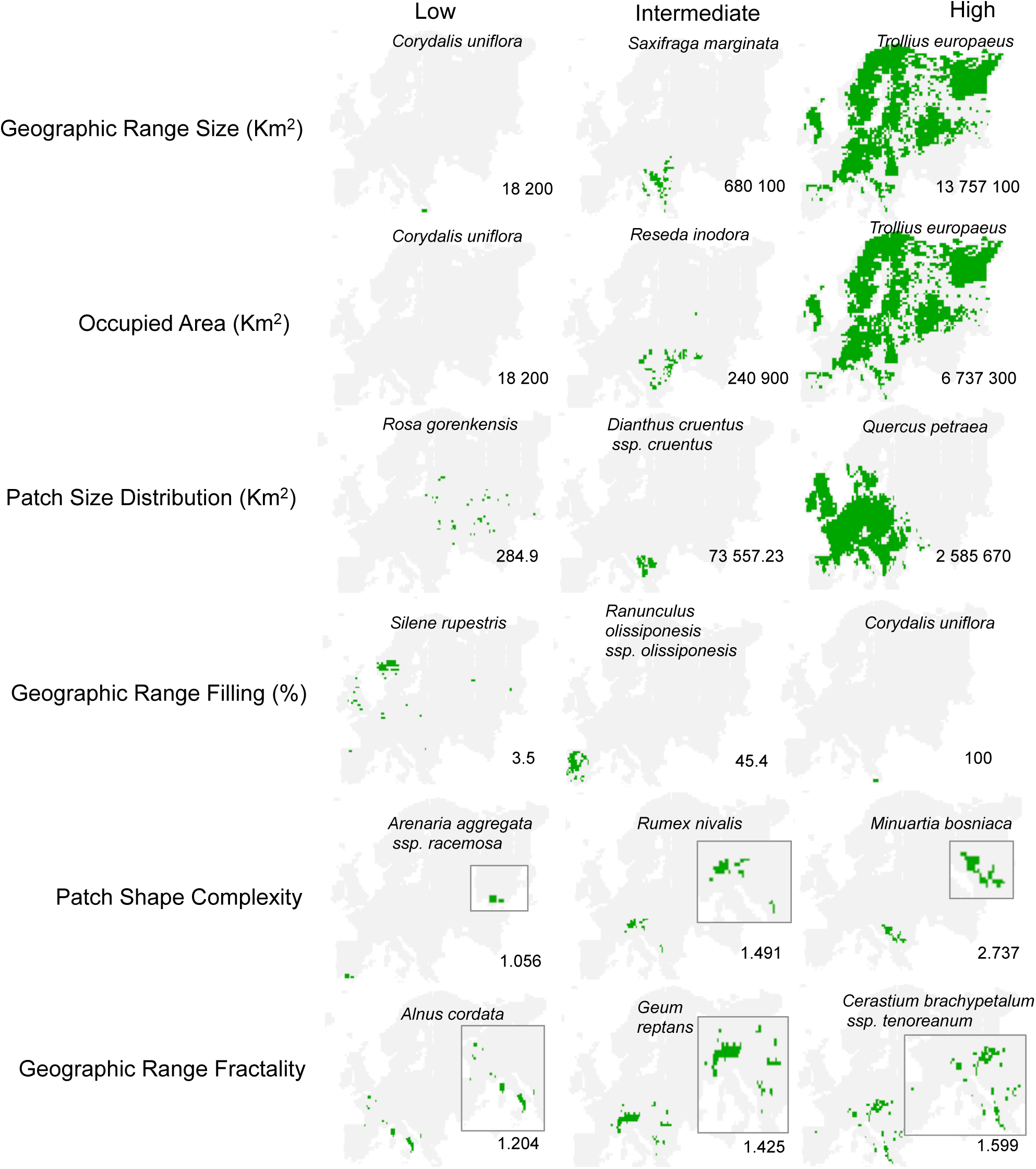
Example species with low, intermediate and high values of geographic range size and structure metrics from a set of 813 European endemic vascular plants, calculated from observed occurrences. Insets represent magnified range maps. Metric values are shown in the bottom right corner of each map.

The geographic range metrics were not sensitive to map resolution, rasterization approach of occurrence data, or approaches taken to estimate species’ geographic range boundaries (Appendix S2.3.1-S2.3.3). For a large number of species, classical SDM projections tended to underestimate geographic range size, occupied area and geographic range filling compared to the ESM projections, and the patch shape metrics calculated from maps produced with the two different techniques were poorly correlated (Appendix S2.3.4). Geographic range metrics based on binary maps produced by different thresholding methods were highly correlated, with few exceptions (Appendix S2.3.5). All results presented in the main text are thus based on binary maps thresholded using MaxTSS. The area metrics and the range division metrics decreased, and the patch shape metrics increased significantly with increasing habitat suitability threshold (p<0.05) (Appendix S2.3.6), indicating lower availability and higher irregularity of highly suitable habitat patches within species’ ranges.

### Differences between observed and predicted range metrics

The phylogenetically uncorrected paired t-tests indicated that mean geographic range size, occupied area, patch size distribution and geographic range filling of observed ranges were significantly lower than those based on predicted suitable ranges (|t| > 5.90, p<0.001). In contrast, the phylogenetically corrected paired t-test indicated that only geographic range filling was significantly different, and lower, for observed compared to predicted ranges (|t| = 5.93, p < 0.001) (Fig. 3, Appendix S4.1). Accordingly, tests of phylogenetic signal in each of the range structure metrics separately indicated significant evolutionary relationships between species in observed and predicted geographic range size, occupied area and patch size distribution (Pagel’s λ > 0.731, p <0.001) (Appendix S4.2). Therefore, differences between observed and predicted values for these three metrics were likely structured by evolutionary relatedness between species.

**Figure 3.**
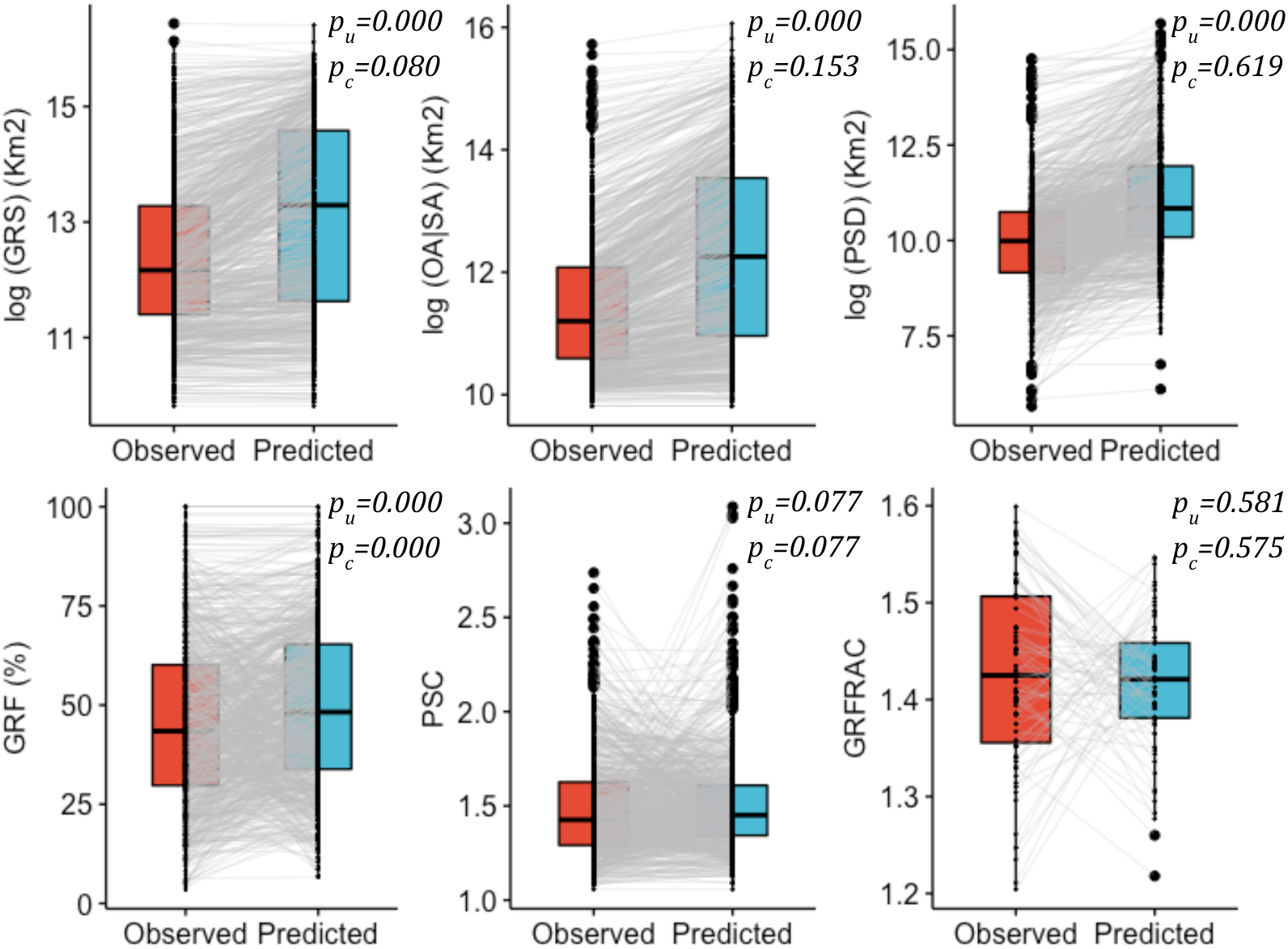
Comparisons of geographic range size and structure metrics quantified from observed and predicted distribution maps for European plant endemics. Significance values of phylogenetically uncorrected (p_u_) and corrected (p_c_) pairwise *t*-tests are shown on the upper right corner of each panel. Significant differences between observed and predicted suitable metrics for geographic range size (GRS), occupied area|suitable area (OA|SA) and patch size distribution (PSD) were explained by phylogenetic relatedness. Differences between observed and predicted suitable geographic range filling (GRF) remained significant after phylogentic correction. No significant differences were found for patch shape complexity (PSC) and geographic range fractality (GRFRAC) in either test. Sample size = 813 species, except for GRFRAC, where analyses were limited to 71 species. The maximum value of the y axis for GRS and OA|SA nears the area of the European continent: log(10.18 million Km^2^) = 16.136, and the minimum value represents log(18 200 Km^2^) = 9.81.

The log response ratio between observed and predicted metrics differed in magnitude across metrics (Fig. 4, Fig. 5, Appendix S4.3). Median log response ratios closest to zero with the narrowest range, and hence best observed to predicted suitable matches, were for the patch shape metrics (Fig. 4e, f) and the largest medians and ranges of log response ratios, and hence largest divergence between observed and predicted suitable, were detected for the range size metrics and patch size distribution (Fig. 4a, b, c). There were major differences between range size and range structure metrics in the shape of the log response ratio relative to the observed metric. In the range size metrics, higher and largely positive log response ratios were common for medium sized observed values (positive quadratic effects, p < 0.001) (Fig. 4a, b). For the range structure metrics, log response ratios were often higher for low observed values and, conversely, lower for high observed values (negative linear or quadratic effects, p<0.001) (Figure 4c - 4f). Exceptions were extreme high values of patch size distribution and geographic range filling, which showed higher, or no change in log response ratio respectively. With the exception of geographic range fractality (Rsq = 0.75), models of all other metrics explained a moderate proportion of variance in data (Rsq < 0.48).

**Figure 4.**
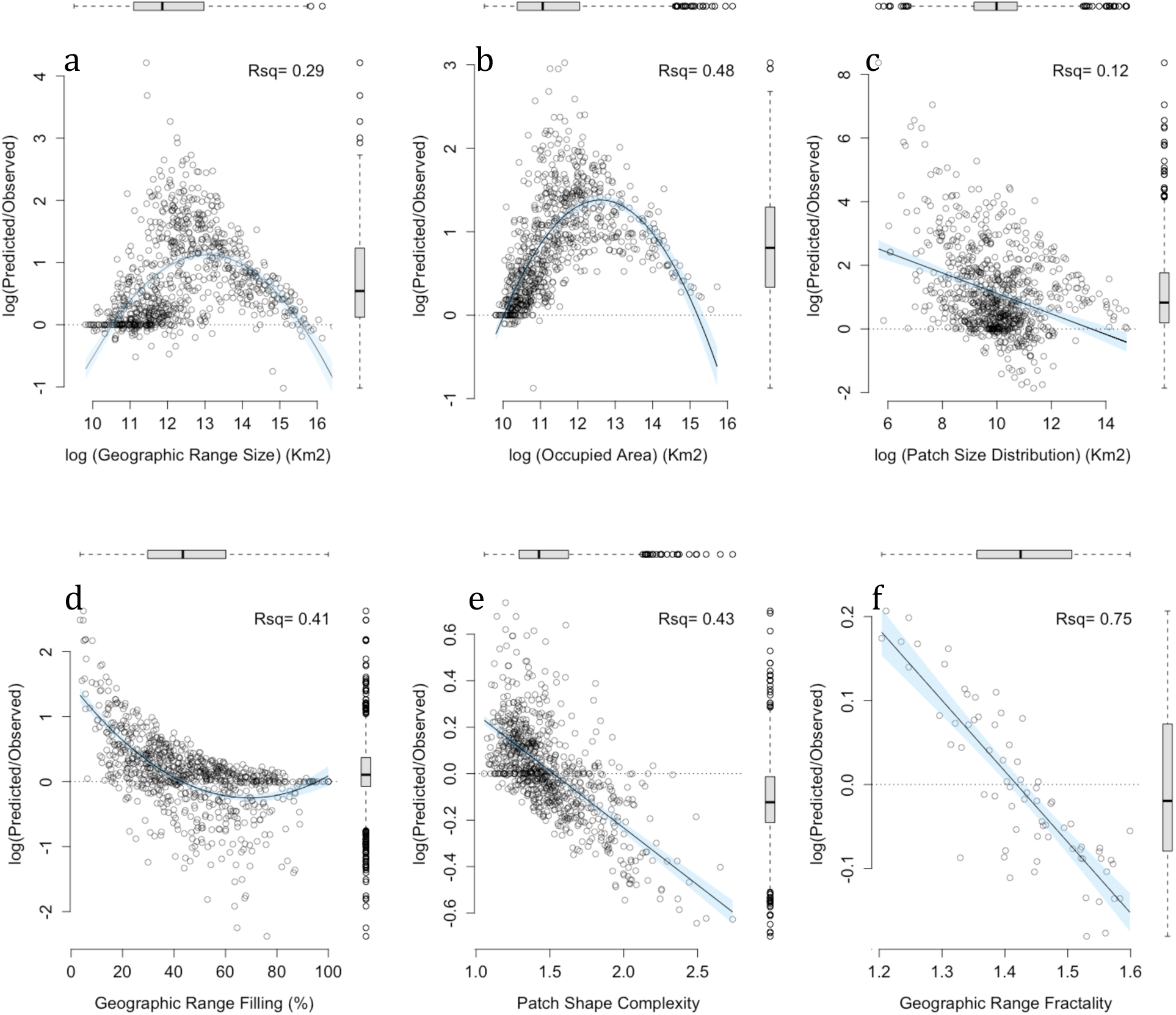
The relationship between the geographic range size and structure metrics quantified using observed occurrence data (x axis) and the log response ratio between the metrics calculated from habitat suitability maps and from observed distributions (y axis). The zero line indicates no difference between the observed and predicted metric. Rsq = R squared value of the linear regression model. The scale of the y axes differs between different metrics. Horizontal and vertical boxplots represent summary values on the x and y axes respectively.

**Figure 5.**
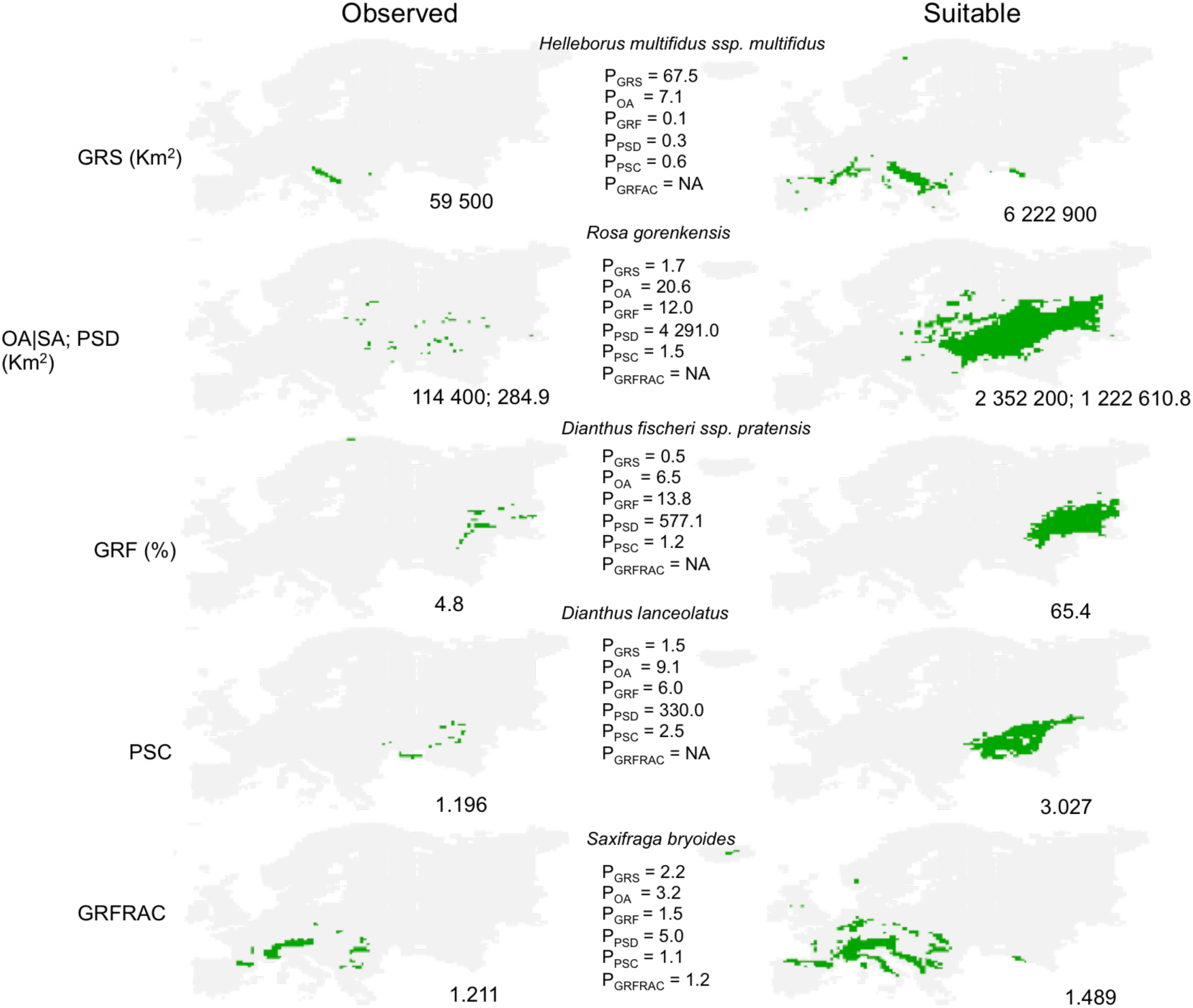
Examples of geographic range size and structure metrics quantified from habitat patches occupied and predicted suitable, and the proportion (P) between the geographic range structure metrics quantified from habitat patches predicted suitable and occupied patches in five example species. Numbers next to the maps indicate the value of the geographic range structure metric. Species were selected to illustrate the largest proportion between predicted and occupied metric (P) from a pool of 813 species endemic to Europe. Proportions of different range structure metrics differ within the same species and between species. Values above 1 indicate larger values of the predicted relative to the observed pattern, values below 1 indicate larger values of the observed relative to the predicted pattern. GRS = geographic range size, OA|SA = occupied (suitable) area, GRF = geographic range filling, PSD = patch size distribution, PSC = patch shape complexity, GRFRAC = geographic range fractality.

### Congruent and divergent drivers of the observed and predicted range metrics

Congruent effects of biophysical and niche variables on observed and predicted range metrics were mostly found for the observed and predicted range size metrics, which were significantly positively correlated with species’ niche breadth and species’ median range latitude. In addition, there was a small significant negative interaction between species’ niche breadth and species’ latitude for both predicted and observed range size metrics. Thus, species’ observed and predicted range size and area increased more rapidly with increasing niche breadth in species of Southern Europe, compared to species of Northern Europe (Fig. 6a, e, Fig. 7a, e, Appendix S4.4, S4.5). Further, both the occupied and suitable area were significantly smaller in species in topographically heterogeneous areas, and both the occupied and suitable area and range size were significantly smaller in species distributed closer to Europe’s coastline (Fig. 6c, d, Fig.7c, d, Appendix S4.4, S4.5).

**Figure 6.**
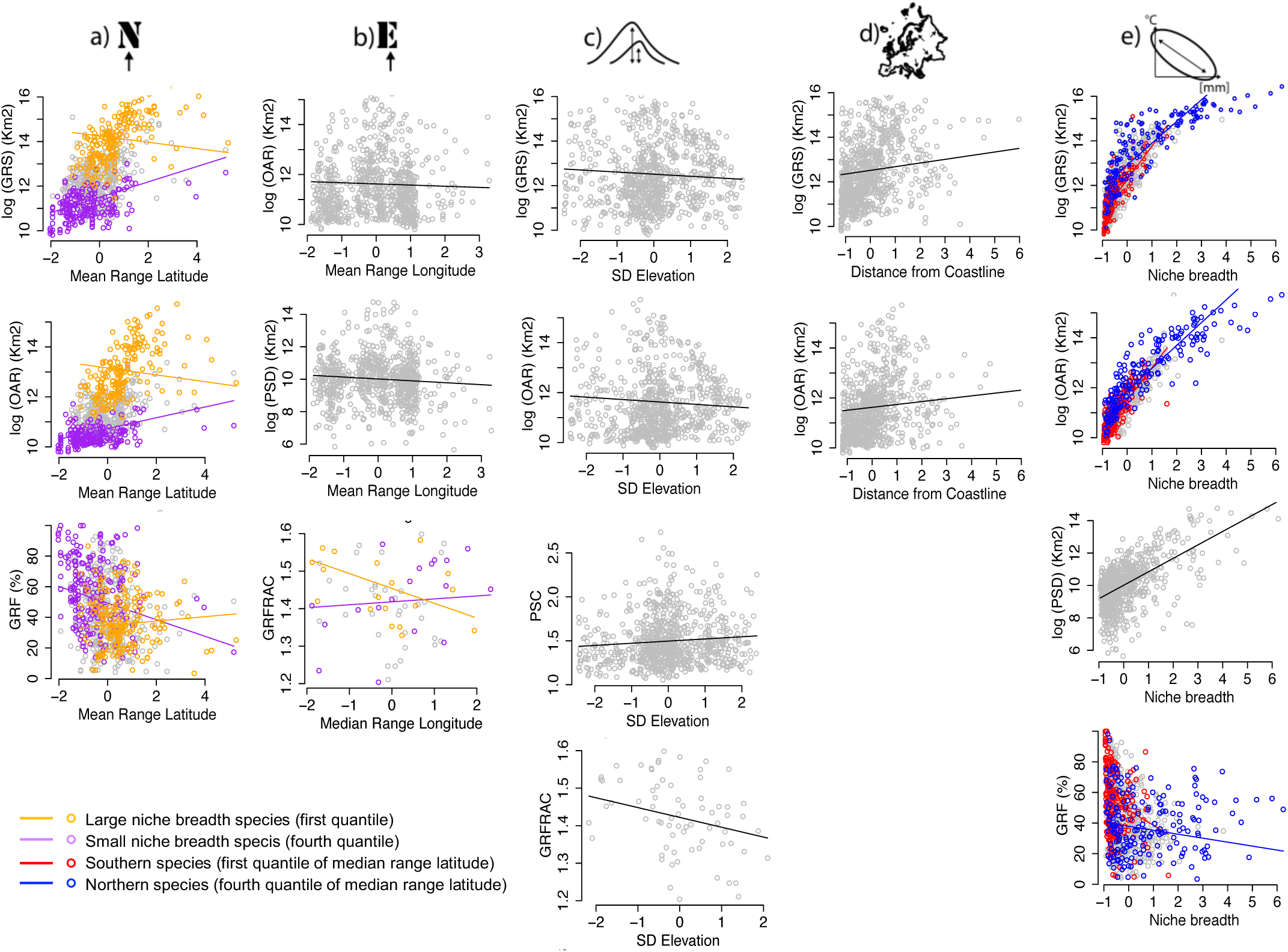
Scatterplots showing the significant effects of **a)** median range latitude, **b)** median range longitude, **c)** topographic heterogeneity (SD elevation), **d)** distance from coastline and **e)** niche breadth on the geographic range structure metrics calculated from observed distributions, modeled using PGLS. GRS=geographic range size, OA=occupied area, PSD=patch size distribution, GRF=geographic range filling, PSC=patch shape complexity, GRFRAC=geographic range fractality. Points represent species, and fitted lines represent the conditional effect of the individual variable, holding other variables constant. Values on the x axes are centered on zero with unit variance. Effect sizes and model details are presented in Supporting material.

**Figure 7.**
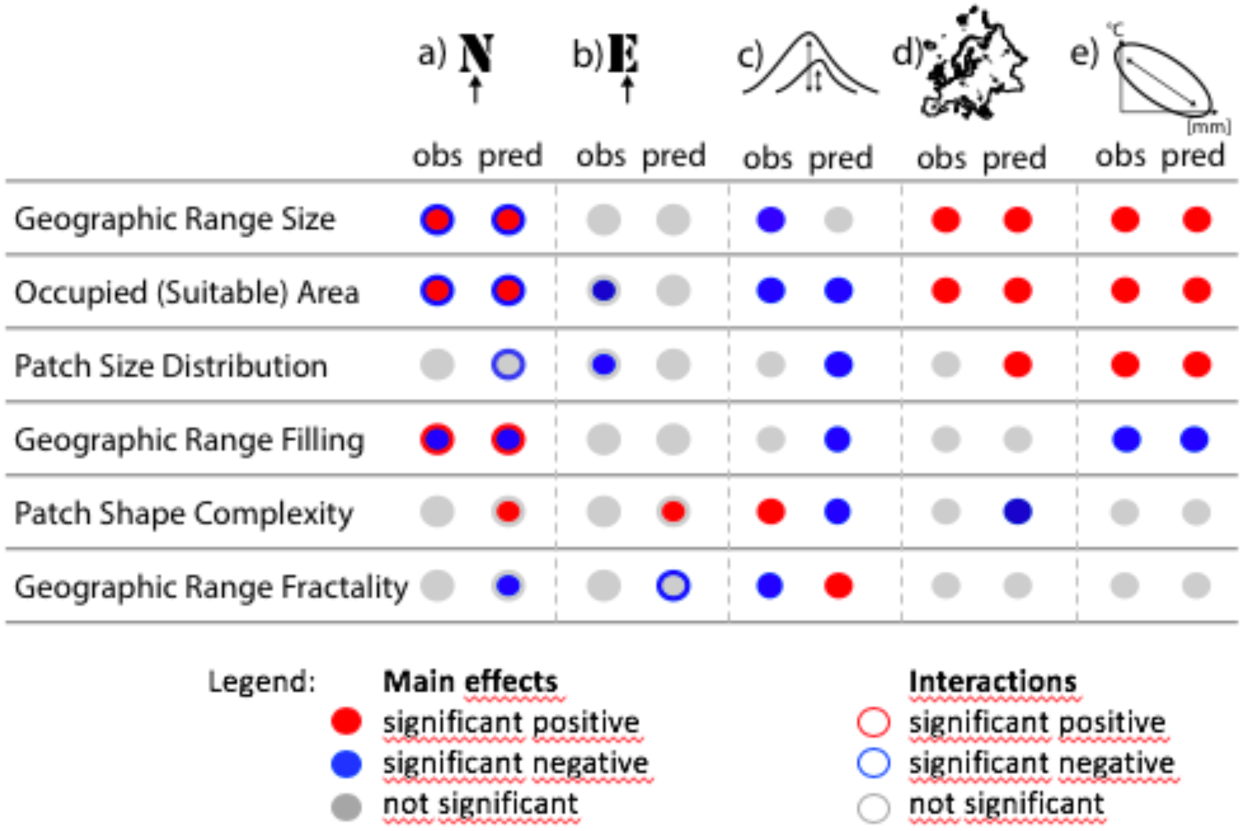
Diagram showing significant effects of **a)** Median range latitude, **b)** Median range longitude, **c)** Topographic heterogeneity (SD Elevation) **d)** Distance from coastline, and **e)** Niche breadth on six geographic range structure metrics modeled using PGLS on species’ observed and predicted distributions. Effect sizes and model details are presented in Figure 6 and Supporting material.

Divergent effects of biophysical and niche variables on observed and predicted range size metrics were found only in two instances: 1) occupied area was significantly lower in species of Eastern Europe, but suitable area was not correlated with median range longitude (Fig. 6b, Fig. 7b, Appendix S4.4, S4.5), and 2) observed range size was significantly lower in species distributed over large elevation gradients, but predicted range size was not related to topographic complexity (Fig. 6c, 7c, Appendix S4.4, S4.5).

Of the range structure metrics, congruent effects were only found for observed and predicted range filling, both significantly negatively correlated with median range latitude and species’ niche breadth, and there was a significant positive interaction of niche breadth and latitude on this metric (Fig. 6a, e, Fig. 7a, e, Appendix S4.4, S4.5). Thus, range filling decreased strongly with increasing niche breadth in Southern Europe, but to a lesser extent in Northern Europe.

Divergent effects of geographic and ecological variables were common between observed and predicted range structure metrics. While for predicted patch size distribution we found a significant negative interaction between niche breadth and latitude, the observed patch size distribution was affected, positively, only by species’ niche breadth (Fig. 6e, Fig. 7a, e, Appendix S4.4, S4.5). Predicted patch size distribution was not correlated with range longitude, but the observed patch size distribution was significantly lower in species of Eastern Europe (Fig. 6b, Fig. 7b, Appendix S4.4, S4.5). Predicted patch size distribution decreased significantly with increasing topographic heterogeneity and nearer to Europe’s coastlines, but the observed patch size distribution was not affected significantly by either parameter (Fig. 7c, d, Appendix S4.4, S4.5).

Predicted geographic range filling was significantly lower in topographically complex areas, but the observed range filling did not vary with topographic heterogeneity (Fig. 7c, Appendix S4.4, S4.5).

Predicted patch shape complexity was significantly higher in species distributed at high latitudes, longitudes and significantly lower near coastlines, but the observed metric did not respond to any of these parameters. Predicted patch shape complexity decreased significantly with increasing topographic heterogeneity, but the opposite effect was found in the observed metric (Fig. 6c, 7c, Appendix S4.4, S4.5).

Predicted geographic range fractality was significantly lower at high latitudes, but the observed metrics did not vary with latitude. Predicted range fractality did not correlate with longitude, but there was a significant negative interaction between niche breadth and longitude for the observed geographic range fractality, hence in species with larger niche breadth, the observed complexity of patch shapes was more even across a range of patch sizes than predicted in species of Eastern Europe (Fig. 7a, b, d, Appendix S4.4, S4.5). Predicted geographic range fractality increased significantly with increasing topographic heterogeneity, but the opposite effect was found in the observed metric (Fig. 6c, 7c, Appendix S4.4, S4.5).

The pGLS models explained a large proportion of variation in observed and predicted geographic range size, occupied (suitable) area and predicted patch size distribution (0.669 < adj. R squared < 0.836), and a lower proportion of variation in observed patch size distribution and observed and predicted geographic range filling, patch shape complexity and geographic range fractality (adj. R squared observed < 0.328 and predicted < 0.186) (Appendix S4.4).

Phylogenetic signal was detected only in pGLS models of occupied area (Model λ: 0.222, Appendix S4.4).

## Discussion

Using complete occurrence data from atlas maps at ~50 km resolution and species distribution models for over 800 plant taxa endemic to Europe, we show that the internal structure of species’ observed ranges varies substantially among species, and cannot be inferred consistently from measures of range size, nor from SDM predictions. Differences between species’ observed and predicted range and area size and patch size distribution are phylogenetically structured. While geographical factors and niche breadth affect the size and structure of species’ ranges, the explanatory variables differ qualitatively and quantitatively between observed and predicted range metrics. Range structure metrics are important to consider independently of, or in addition to, range size metrics when interpreting biogeographic patterns of observed or predicted occupancy.

### Range structure is largely decoupled from range size

Range size metrics, in particular those that incorporate both unoccupied and occupied areas within the range boundaries are not congruent with species’ internal range structure. The negative correlation between range size and range filling supports the view that mechanisms shaping species’ range size (extent of occurrence) and the ability of species to fully occupy it (range occupancy) may differ substantially (Gaston et al. 2003). The observed patch size distribution was better correlated with occupied area than with the observed geographic range size, suggesting that species’ occupied area could be linked to processes that create abundance-occupancy patterns at the landscape scale (Freckleton et al. 2006, Meloni et al. 2017), while species’ range size can additionally be shaped by spatio-temporal processes that generate occurrence patterns, such as long-distance dispersal or spatial rearrangement of populations with geographic range shifts (Hampe and Jump 2011). Analysing which landscape-level processes scale up to structure species’ biogeographic ranges has remained largely unexplored (Kent 2007), yet our results suggest that it may be key to understanding and predicting internal range rearrangements in changing environments.

### Differences between observations and predictions are not consistent among and within range size and structure metrics

Differences between observed and predicted geographic range size, area and patch size distribution were due to phylogenetic relationships between species. One reason for the phylogenetically structured mismatches between these predicted and observed metrics could be a higher range disequilibrium with the environment of particular groups of closely related species, which would lower the ability of SDMs to correctly predict distributions (Guisan et al. 2017). Environmental stability may promote phylogenetic conservatism and it also modulates species’ range size (Morueta-Holme et al. 2013, Zacaï et al. 2017). We hypothesize that range disequilibrium due to large-scale climatic events on the continent (e.g. glaciations) or human land use limiting dispersal (Miller and McGill 2017) were more dramatic in selected groups of phylogenetically related species, while species with closer observed to predicted matches benefited from more stable climates. Another hypothesis posits that phylogenetically older species may have more fragmented ranges than their younger relatives (Sheth et al. 2020), due to e.g., increasing complexity of biotic interactions with species’ age. While these hypotheses need further testing, our results underscore that we lack a good understanding of the phylogenetic structuring of species’ ranges (Gaston et al. 2003, Sheth et al. 2020).

The only metric with higher predicted relative to observed values, independently of the phylogenetic relationships between species, was geographic range filling. Consequently, it is very common that SDMs extend species’ range boundaries with remote suitable habitat patches that are likely to fall beyond species’ normally observed colonisation abilities (Estrada et al. 2015).

Different sets of species were responsible for the divergence between observed and predicted range size and range structure metrics. For the range size metrics, inferring realized from predicted range and area sizes was the most problematic for species with intermediate observed values of these metrics. This may arise due to both methodological and ecological constraints: for small ranges, the convergence between observations and predictions may be a modeling artefact due to models being more easily overfit; for large ranges, the convergence might be due to geographical constraints from landmass availability and continental barriers, which can drive species’ global range size especially in highly dispersive species (Broennimann et al. 2006, Sheth et al. 2020). Alternatively, both narrow climate endemics of Europe (Ohlemüller et al. 2008) and species with large niches might be less prone to modeling biases from niche truncation.

For the range structure metrics, inferring realized from predicted patch size distribution and range filling was problematic in species with the smallest observed values of these metrics. In such species with constrained range occupancy - but not necessarily small range sizes -, overpredictions from SDMs due to missing limiting variables (dispersal limitation, biotic interactions, human impact, etc.) are highly likely. Consequently, the predicted patch size distributions and spatial arrangement of suitable habitat patches within species’ ranges are not realized, or conversely, are eroded, and theoretical inferences about the associated spatial processes need careful testing.

Discrepancies between predicted and observed patch shapes raises modeling difficulties of a different nature. The predicted simplification of the complex observed shapes, which as a result modulates discrepancies between predicted and observed range fractality, suggests that SDMs are blurring patch boundaries (also indicated by the lower variance of predicted than of the observed patch shape metrics in Fig. 3). Distribution borders theoretically self-arrange along complex geometrical patterns (Oborny 2018) that are very likely lost during the predictive modelling process and over coarse spatial resolutions.

These important divergences between metrics and between species severely complicate current and future predictions of species’ internal range rearrangement under climate change, and we cannot hereafter ignore the direction and magnitude of the differences when we infer realized distributions from predicted habitat suitability maps.

### Divergent responses of observed and predicted range size and structure to geographical and ecological factors

Species’ niche breadth was a strong biogeographic determinant of species’ ranges, with similar effects on the predicted and observed range metrics. The widely recognized niche effects on range and area size (Moore et al. 2018, Sheth et al. 2020) were associated with further niche effects on species’ range structure (observed and predicted geographic range filling and patch size distribution, and predicted range fractality). Niche breadth can therefore be a useful tool in global predictions of species’ range structure and internal range rearrangements, especially in a climate change context (Broennimann et al. 2006, Connor et al. 2018). However, niche effects on range structure metrics were modulated by range latitude and longitude, and the relatively homogeneous climate over large areas in Northern Europe (Ohlemüller et al. 2008) might have enabled species with different niche breadths, that include the Northern European climate, to acquire relatively large range and area sizes, filled with large suitable patches. This result contradicts the view that towards higher latitudes larger range sizes are due solely to species’ increasing ecological tolerance (Rapoport’s rule:, Rapoport 1982, Stevens 1989). Ecological tolerance does not seem to exclusively govern geographic ranges (Gaston and Chown 2016, Payne and Smith 2017, Moore et al. 2018), instead it might modulate species’ range size and structure within the spatial availability of geophysical conditions, according to the principles of Hutchinson’s niche-biotop duality (Colwell and Rangel 2009).

Constraints on species’ predicted and observed area and range sizes in Southern Europe, in topographically heterogeneous areas, as well as near the coastlines were often, but not always, associated with additional constraints on range structure. The historical imprints of the higher climate stability in Southern Europe during glaciations, associated with a diversity of small climate pockets (Ohlemüller et al. 2008), the higher rates of speciation in topographically complex areas (Morueta-Holme et al. 2013, Steinbauer et al. 2016), and the geographical barriers imposed by the continental coastlines (Broennimann et al. 2006) on species ranges could therefore also be manifest in species’ internal range geometry, with potentially important ecological and evolutionary consequences that need further exploration.

However, most effects of geographical parameters on the predicted range metrics were not transferable to the observed metrics, and we suspect biological and historical, as well as methodological reasons underlying these inconsistencies. The relaxed dependency of predicted patch size distribution on niche breadth towards the North, undetected in observed patch size distributions, could be due to at least two factors: 1) patch size distribution is modulated by species’ biological properties that manifest regardless of species’ range position, such as dispersal ability (Freckleton et al. 2006, Pagel et al. 2019) and 2) there is a strong postglacial dispersal lag in species distributed in Northern Europe, which leaves large parts of suitable habitat patches unoccupied (Svenning et al. 2006). Historical factors or anthropic pressures might have similarly reshuffled, or eroded, theoretical geographical patterns of Eastern Europe (Kajtoch et al. 2016), where we revealed an observed, but not predicted, decline in area and patch size distribution, as well as predicted, but not observed, increase in patch shape complexity. Topography and geographical barriers were not enough to explain realized patch size patterns either, because small observed patch size distributions due to small average patch size or small overall area were not only frequent in highlands and near the coastlines as expected, but also in the more homogenous lowlands and inland. Thus, human land use, colonization and fragmentation histories, or species’ intrinsic properties, are most likely modifying the predicted theoretical internal range geometries, obscuring the accuracy of future range forecasts (Svenning et al. 2006, Estrada et al. 2014, Miller and McGill 2017).

### Limitations and future directions

Due to the coarse resolution of our dataset, we only tested drivers of range size and structure that may operate over large biogeographic scales, such as niche breadth, phylogeny, coarse scale geomorphology, large geographic barriers and coarse geographic gradients. The repertoire of metrics that describe species’ range size and structure structure can definitely be expanded in the future and approaches can certainly be refined (Sheth et al. 2020). Some of the metrics chosen in this study were weakly responsive to the chosen geophysical and ecological drivers but may better respond to other factors yet to be tested. Smaller effect sizes with large confidence intervals for predictions of patch shape complexity and range fractality metrics emphasised that these metrics are poorer predictors of processes at global scales, similarly to findings of Pearson et al. (2014). Coarse map resolutions can affect global range predictions (Connor et al. 2018), and the smoothing of the range shapes by the distribution models may become problematic when aiming for accurate projections of species’ internal range geometry into the future. Further efforts should be made to identify which aspects of species’ biology or of the geophysical space are linked to biological patterns at given spatial resolutions (Connor et al. 2018, Pagel et al. 2019), and to improve species distribution models with finer predictors able to better reflect the peculiarities of species’ internal range structure.

## Conclusions

Our results have important implications for the development and interpretation of predictive models of species expansion, persistence and extinction in response to climate change. Fragmentation of occupied area or suitable habitat has already been identified as a better predictor of extinction risk than range size (Pearson et al. 2014, Crooks et al. 2017), and we propose that factors influencing different aspects of species’ range structure, in addition to those that influence range size, may be useful conservation targets (as in Cianfrani et al. 2018). The predictable inconsistencies between observed and current predicted range structure metrics and their drivers enable us to better approximate uncertainties in species predictions. Some of the discrepancies might stem from species’ inability to occupy all habitats predicted suitable, while others may be linked to our inability to capture accurately species’ responses to environmental constraints when modeling distributions. By examining which species are sensitive to different properties of species’ ranges, and the sensitivity of populations to changing range configurations, we can refocus conservation efforts to search for critical range structure thresholds for species at the highest risk.

## Acknowledgements

We thank AFE for distribution data. We thank Carsten Hobohm for kindly providing the list of European endemic species in EvaplantE, the database on endemic vascular plants in Europe. We thank Ruben Mateo for help with CGRS grid calculations in early stages of the project and Martha Liliana-Serrano and Liam Revell for readily helping with phylogenetic analyses. We thank Beáta Oborny for very useful comments on manuscript versions in the making. AMC was funded by the Marie Sklodowska-Curie Individual Fellowship GEODEM-658651 under the EU Horizon 2020 Framework Programme for Research and Innovation of the European Research Council. YMB was supported by an Irish Research Council Laureate Award IRCLA/2017/60. AG was supported by a Swiss National Science Foundation grant (CR23I2_162754).

## Biosketch

Anna M. Csergő led this work as a Marie Sklodowska-Curie postdoctoral research fellow at Trinity College Dublin, Ireland and then as associate professor at Szent István University, Hungary (https://annamariacsergo.weebly.com/). Together with the co-authors she is interested in how species’ life history and population-level processes interact with the geometry and suitability of habitat fragments within a species’ range to modulate global persistence patterns.

## Data accessibility statement

Data associated with this manuscript will be made openly available in the Dryad data repository upon acceptance of the manuscript.

