## Supporting material for "Beyond range size: drivers of species’ geographic range structure in European plants"

**Appendix**

**S1. Species distribution modeling**

*Environmental data selection.* To characterise environmental and geographic variation throughout species’ ranges, 20 climate, 7 soil and 2 topographic raster layers were used initially. Climate variables were downloaded from the CliMond Archive (V1.2; https://www.climond.org/BioclimRegistry.aspx, Kriticos et al. 2012) and the following variables were chosen: Bio01, Bio02, Bio03, Bio04, Bio05, Bio06, Bio07, Bio08, Bio09, Bio10, Bio11, Bio12, Bio13, Bio14, Bio15, Bio16, Bio17, Bio18, Bio19, Bio28 (Kriticos et al. 2014). Soil physical and chemical properties (bulk density, field capacity, profile available water capacity, soil carbon density, total nitrogen density, soil thermal capacity, wilting point) were sourced from the International Geosphere-Biosphere Programme IGBP (Global Soil Data Task 2014), and topographic heterogeneity metrics (aspect, slope) were calculated from elevation maps (Jarvis et al. 2008) using the ‘raster’ package in R (Hijmans and van Etten 2012). Topographic aspect was distance from North expressed in degrees, where North represents 0 degrees (Roberts 1986). All layers were downloaded at 30’’ resolution (~1✕1 km^2^) except soil data which was downloaded at 5’ resolution (~10✕10 km^2^) and subsequently resampled to 30’’.

AFE maps use the Military Grid Reference System (MGRS), an extension of the UTM system. To extract environmental data for each AFE grid cell, we used the Common European Chorological Grid Reference System (CGRS), a modified MGRS, with a resolution of ~50✕50 km^2^, downloaded from European Environment Agency portal on 2015 June 29 (http://www.eea.europa.eu/data-and-maps/data/common-european-chorological-grid-reference-system-cgrs). We calculated mean, minimum and maximum across the 1✕1 km^2^ pixels contained in each 50✕50 km^2^ grid cell.

To avoid overparameterization and reduce correlations among variables (Harrell et al. 1984), we performed a cluster analysis based on Unweighted Pair Group Method with Arithmetic Mean method (UPGMA) on a correlation matrix for mean, minimum and maximum values previously calculated for the climate, soil and topographic variables. The resulting tree was pruned at 0.5 correlation value, and one variable per cluster was selected, with the exception of the largest cluster, from where two variables correlated at 0.7 cutoff value were selected. Selection of candidate variables within each cluster was based on visual inspection of their spatial variability across Europe. This selection approach resulted in 14 final predictors: five predictors related to temperature: tdr.max (mean diurnal temperature range, Bio02, maximum), tar.min (temperature annual range, Bio07, minimum), twetq.min (mean temperature of wettest quarter, Bio08, minimum), tdryq.max (mean temperature of driest quarter, Bio09, maximum), twarmq.max (mean temperature of warmest quarter, Bio10, maximum), three were related to precipitation: p.mean (annual precipitation, Bio 12, mean), ps.max (precipitation seasonality, Bio15, maximum), pdryq.min (precipitation of driest quarter, Bio17, minimum), five were related to soil properties: totaln.min (total nitrogen density, minimum), totaln.max (total nitrogen density, maximum), thermcap.mean (thermal capacity, mean), wiltpont.min (wilting point, minimum), wiltpont.max (wilting point, maximum), and one was related to topography: slope.max (slope, maximum) (Figure S1.1).

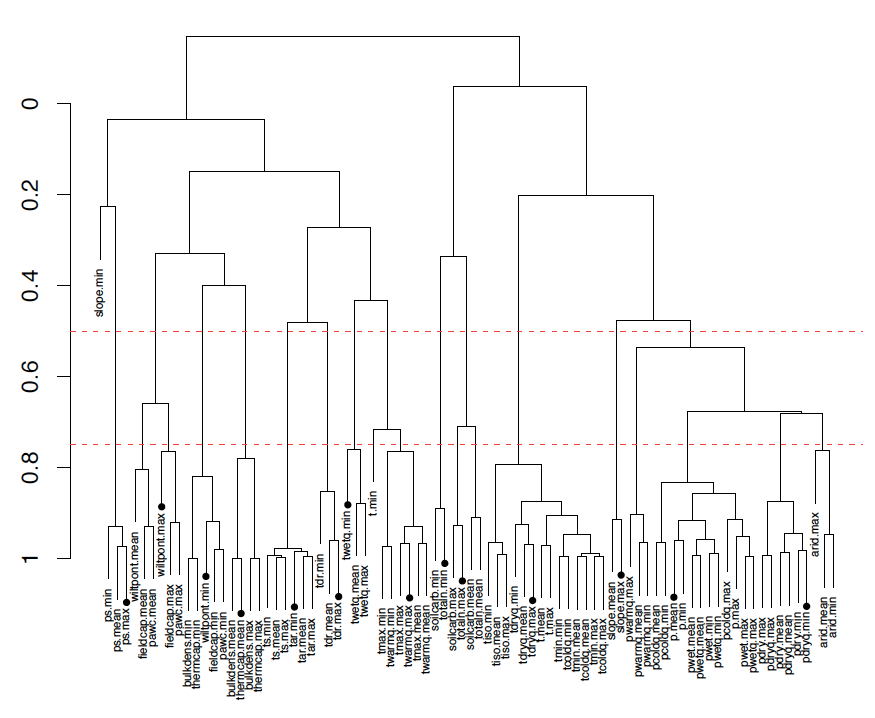

**Figure S 1.1.** UPGMA dendrogram showing the correlation between all environmental variables considered and the position of final selected variables, indicated with black dots. The cutoff correlation values of 0.5 and 0.7 respectively are marked by the dotted red lines.

*Modelling approaches*

*Ensemble of Small Models (ESM)*

We built Ensemble of Small Models (ESM; Breiner et al. 2015) to predict the habitat suitability maps for each species using the 14 environmental variables previously selected. With this technique, bivariate models are first fitted with two explanatory variables at a time, resulting in the fitting of bivariate models with all possible pairs of environmental predictors (i.e. 14x13/2=84). Bivariate models were fitted respectively using Generalized Linear Models (GLM, McCullagh and Nelder 1989), Random Forest (RF, Breiman 2001) and Maxent (Phillips and Dudik 2008). A total of 1365 models were fitted for each species (3 techniques x 84 combinations x 5 iterations). All grid cells from which species were not reported were considered real absences, and a weight was applied in the calibration of the models so that presences and absences have equal (0.5) prevalence. Models were calibrated with five random subsets of 70% of the data and evaluated with the remaining 30% of the datasets. We used the maximum value of TSS (Allouche et al. 2006) across all possible probability thresholds (between 0 and 1) to assess model performance (maxTSS; Guisan et al. 2017). The final ensemble model was constructed from all models taking a weighted average of the predicted occurrence probabilities proportional to the maxTSS values. The mean and standard deviation of maxTSS were calculated across all runs and techniques. Because the AFE cells are not homogeneous in area and shape, we rasterized the observed and predicted distributions to 10 km.

*Classical Species Distribution Models (SDM)*

In addition to our main approach (Ensemble of Small Models, ESM; Breiner et al. 2015) we performed classical species distribution models based on ensemble technique (SDM; Thuiller et al. 2009). The SDMs were performed using ensemble modeling as implemented in the BIOMOD2 library (Thuiller et al. 2009). For this approach we selected species with at least 10 occurrences. As different statistical techniques can produce different results (Elith et al. 2006), SDM models were run with three different techniques: generalized linear models (GLM, McCullagh and Nelder 1989), random forest (RF, Breiman 2001) and maximum entropy modeling (MAXENT; Phillips and Dudik 2008) with no interaction between terms and all other parameters set to their default values as implemented in BIOMOD2. All grid cells from which species were not reported were considered real absences, and a weight was applied in the calibration of the models so that presences and absences have equal (0.5) prevalence. Models were calibrated with five random subsets of 70% of the data and evaluated with the remaining 30% of the datasets. We used TSS (Allouche et al. 2006) to assess model performance. The final ensemble model was constructed from all models, taking a weighted average of the predicted occurrence probabilities proportional to the TSS values. The mean and standard deviation of TSS were calculated across all runs and techniques.

Both species distribution modeling methods had generally high predictive power with high values of the TSS evaluation metric: 0.887/0.926/0.992 for ESMs and 0.973/0.984/1 for SDMs  (1^st^ Quartile/Mean/3^rd^ Quartile) (Figure S1.6).

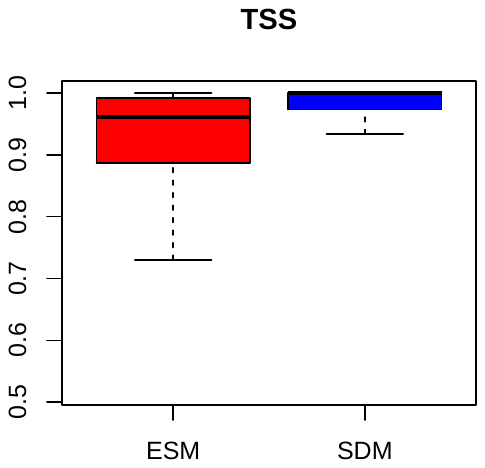

**Figure S1.2** Boxplots showing the TSS evaluation metric values for two types of species distribution models: Ensemble of Small Models (ESM) and conventional Species Distribution Models (SDM) (N = 827 species).

**S2 Calculation of geographic range metrics, relationships between range metrics and sensitivity analyses**

**S2.1 Calculation of geographic range metrics**

To calculate geographic range metrics we used landscape metrics developed in Fragstats (McGarigal et al. 2012) adapted to R in the *ClassStat* function of the ‘SDMTools’ package (VanderWal et al. 2015).

We present here definitions of metrics developed for landscape ecology studies in Fragstats, which we adapt to analyses of the size and structure of species’ geographic ranges. Some metrics were renamed or modified to accommodate the *ClassStat* function to geographic range-level interpretation. The metrics were computed from distribution maps in raster format and the definition of terms presented here refers to raster cells that are occupied by a species (occupied habitat) and background or matrix cells that are not occupied (unoccupied habitat). For the predicted maps, the metrics were computed from raster cells representing habitats predicted suitable and background cells representing habitats predicted unsuitable.

**A. Area metrics**

Area metrics inform on the size of species’ geographic range and the size of occupied or suitable habitats within the species’ geographic range boundaries.

**1. Geographic Range Size (km^2^) (GRS):** area of all occupied non-occupied habitat patches within a species’ geographic range boundaries. We defined the geographic range boundaries using a minimum convex polygon around occupied habitat grid cells. The minimum convex polygon was defined around 95% of the occupied grid cells to avoid range size overestimation due to highly disjunct habitats (as in Figure 2.1.1; see section 2.3.3 of this appendix for a sensitivity analysis of the geographic range metrics to this approach).

*GRS = cell size^2^ * n. cells*

, where

*cell size* = raster grid cell size (km)

*n.cells* = total number of occupied and unoccupied grid cells within the species’ geographic range boundaries

*Range:* GRS > 0. The metric approaches 0 when the number of grid cells within the species’ geographic range boundaries is very low.

Note: This metric is called “*Total landscape area (A)*” that includes internal background in Fragstats, and “*L.area*” in SDMTools.

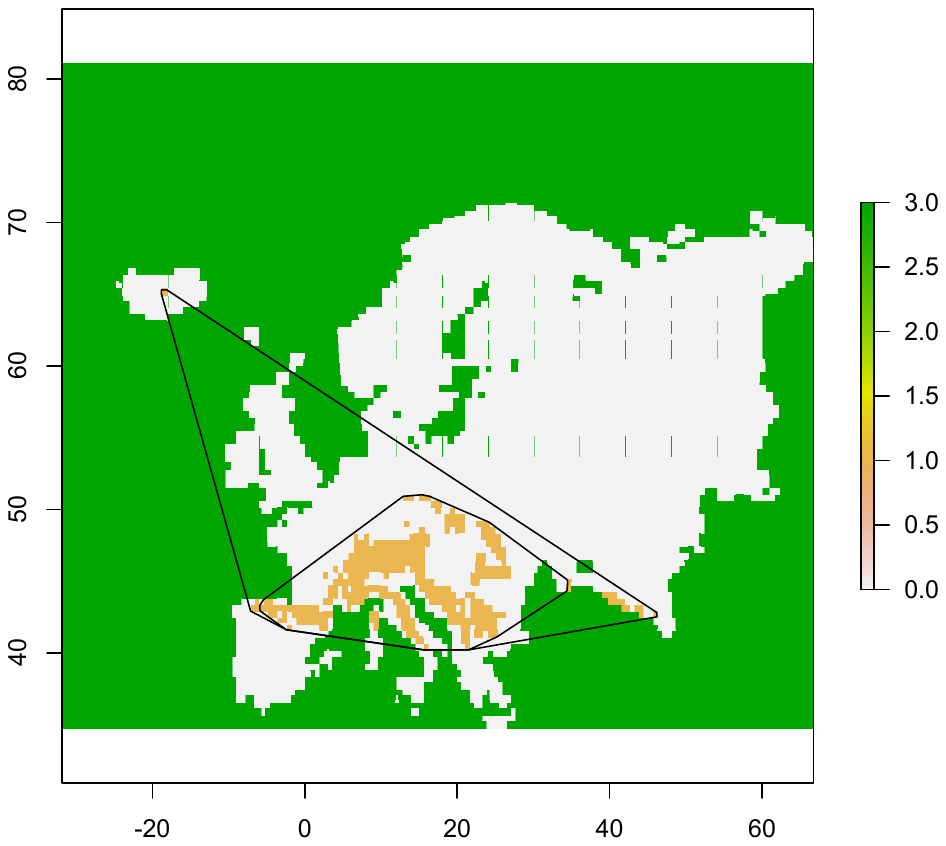
**Figure S2.1.1** Minimum convex polygon defined around 100% of points (outer polygon) and 95% of the points (inner polygon) in an example species.

**2. Occupied Area (km^2^) (Occupied Area, OA):** the sum of the area of all occupied habitat patches within a species’ range (as **GRS** but includes only occupied grid cells within the species’ range boundaries). **Suitable Area (SA)**, when the metric is calculated from grid cells representing suitable habitat

$OA(or SA)=\sum_{i=1}^{n} a_{i}=\sum_{i=1}^{n} {{cell size}^{2}*n. cells}_{i}$

, where

*n*=number of occupied habitat patches within a species’ range

*a_i_* = area of patch *i*

*cell size* = raster grid cell size (km)

*n.cells_i_* = number of cells in patch *i*

*Range:* OA (SA) > 0. The metric approaches 0 when the number of occupied grid cells is very low.

*Note:* The metric is called “*Total Area (TA)*” in Frastats and “*total.area*” in SDMTools. In contrast to Fragstats and SDMTools, we don’t convert area to hectares and the unit of the metric is km^2^ not m^2^.

**B. Range division metrics**

Range division metrics are quantitative measures of the size and spatial arrangement of occupied habitat patches within the species’ geographic range boundaries, relative to species’ geographic range size.

**1. Patch Size Distribution (PSD) (km^2^):** measures the subdivision of a species’ geographic range into occupied habitat patches of different sizes relative to species’ geographic range size. The metric equals the occupied patch area squared, summed across all occupied patches, divided by the geographic range size. It denotes the size of the areas obtained by dividing the species’ geographic range into S parts of equal size that have same degree of geographic range division as obtained for the observed cumulative area distribution (Cumulative area distribution is calculated by regressing the cumulative area of the occupied habitat patches (axis y) against patch size (axis x).) (Jaeger 2000). The metric was proposed under the name “*Effective mesh size*” by Jaeger (2000), and in contrast to the other landscape subdivision metrics proposed in the same work, this metric is area-proportionately additive i.e., it characterizes the subdivision of a species’ geographic range independently of its size, making it suitable for comparative analyses.

$PSD=\frac{\sum_{i=1}^{n} a_{i}^{2}}{GRS}= \frac{\sum_{i=1}^{n} {{cell size}^{2}*n. cells}_{i}}{{cell size}^{2} * n. cells}$,

,where

*n* = number of occupied habitat patches within a species’ range

*a_i_* = area of patch i

*GRS*=Geographic Range Size

*cell size* = raster grid cell size (km)

*n.cells_i_* = number of cells in patch i

*n.cells* = total number of cells within the geographic range boundaries (both occupied and not occupied)

*Range:* The metric is minimum when the area of occupied habitat patches is very small relative to the geographic range size, and it is maximum when one occupied patch fills out the species’ entire geographic range.

*Note 1:* The metric is called **“***Effective mesh size***”** in Fragstats and “*effective.mesh.size*” in SDMTools.

*Note 2:* When comparing the range subdivision across different species, small mesh size can indicate a large number of small patches dispersed throughout a large area of distribution or one patch that fills a small geographic range. Consequently this metric doesn’t differentiate between different types of rarity in Rabinowitz (1981), but it does indicate rarity. The metric is also useful to compare temporal or spatial changes in range subdivision of the same species.

**2. Geographic Range Filling (GRF) (%)**: represents the proportion of a species’ geographic range represented by occupied habitat patches. It equals the occupied area (OA) divided by the geographic range size (GRS), multiplied by 100 (to convert to a percentage).

***GRF = (OA/GRS)*100***

*Range:* 0 < GRF ≤100. The metric approaches 0 when the number of occupied grid cells within the species’ geographic range is low and it equals 100 when the entire geographic range is comprised of occupied grid cells.

*Note:* The metric is called “*Percentage of Landscape*” in Fragstats and ‘*prop.landscape*’ in SDMTools.

**C. Patch shape metrics**

Patch shape metrics inform on the shape complexity of the occupied habitat patches throughout a species’ geographic range.

**1. Patch Shape Complexity (PSC) (without unit):** quantifies the shape complexity of the occupied habitat patches compared to a standard shape (square) of the same size for each patch. By comparing patch shapes to a standard shape, the metric alleviates the dependency of the perimeter/area ratio on patch size, in contrast to the simple perimeter/area ratio metric used in landscape ecology, making it more suitable for comparative range structure analyses. The metric equals the patch perimeter divided by the square root of patch area, multiplied by a constant to adjust for a square standard, averaged over all occupied habitat patches. This index emerged from a diversity index proposed by Patton (1975). The metric excludes any non-occupied cells.

$PSC= \sum_{i=1}^{n} \frac{0.25*p_{i}}{\surd a_{i}}=\sum_{i=1}^{n} \frac{0.25*cell size* {number of perimeter edges}_{i}}{\surd{{cell size}^{2}*n. cells}_{i}}$,

,where

*n* = number of occupied habitat patches within a species’ range

*p_i_* = perimeter of patch i

*a_i_* = area of patch i

*cell size* = raster grid cell size (km)

number of perimeter edges _i_ = number of cell edges that make the perimeter of patch i

*n.cells_i_* = number of cells in patch i

*Range:* PSC ≥ 1. The metric equals 1 when each patch within a species’ range is square and it increases without limit as patch shapes become more irregular

*Note:* This metric is called “Shape index” for single patches in Fragstats and “mean.shape.index” in SDMTools, which averages the “shape.index” across all individual habitat patches.

Note2: We replaced the PSC metric function as it presented in SDMTools:

*tout$mean.shape.index = mean(out.patch$shape.index, na.rm = T)*

with a more straightforward function that follows exactly the description given in Fragstats and which we followed:

*tout$mean.shape.index.new = round(mean(((0.25*out.patch$perimeter)/sqrt(out.patch$area)), na.rm = T),digits=1)*

**2. Geographic Range Fractality (GRF) (without unit):** quantifies how the shape complexity of habitat patches changes across a range of patch sizes i.e. spatial scales. In contrast to simple perimeter-area metrics based on average patch shape such as the mean path shape, it regresses the patch area against patch perimeter with habitat patch as unit of observation (Burrough 1986). It equals 2 divided by the slope of regression line obtained by regressing the logarithm of patch area (km2) against the logarithm of patch perimeter (km). GRF excludes any non-occupied habitat cells.

$$PAFRAC=\frac{2}{\frac{[n\sum_{i=1}^{n} (\ln p_{i}* \ln a_{i})]-[(\sum_{i=1}^{n} \ln p_{i})*(\sum_{i=1}^{n} \ln a_{i})]}{(n\sum_{i=1}^{n} \ln p_{i}^{2})-{(\sum_{i=1}^{n} \ln p)}^{2}}}$$

,where

*n* = number of occupied habitat patches within a species’ range

*p_i_* = perimeter of patch i

*a_i_* = area of patch i

*Range:* 1≤ GRF ≤ 2 The metric takes low values if small and large patches alike have similar patch shape complexity and high values when patch shape complexity increases with patch area. Consequently, low values indicate repeated, self-similar patterns across all patch sizes and could reflect the fact that processes at fine scale are propagated to broad geographic scales (Milne 1988). Conversely, high values could indicate that processes at fine scale (small patches) differ from those operating at larger scale (larger patches).

Note 1: The metric is called “*Perimeter-Area Fractal Dimension*” in Fragstats and “*perimeter.area.frac.dim*” in SDMTools.

Note 2: We replaced the function in SDMTools

*tout$perimeter.area.frac.dim = 2/(((tout$n.patches *sum(log(out.patch$perimeter) + log(out.patch$area))) - tout$total.edge * tout$total.area))/(tout$n.patches * sum(log(out.patch$perimeter^2)) - tout$total.edge^2))*

with the more straightforward approach:

*m <- lm (log(out.patch$area) ~log(out.patch$perimeter))*

*tout$perimeter.area.frac.dim = round(2/as.numeric(coef(m)[2]), digits=1)*

Note 3: Due to its dependence on regression analysis, this metric is more reliable for geographic ranges with a large number of patches that differ in size and for linear perimeter-area relationships. We limited the analysis to species with a minimum of 10 habitat patches. The linearity of perimeter-area relationship should be tested separately.

**S2.2 The relationship between different geographic range metrics**

**Figure S2.2.1** Scatterplots showing the relationship between six geographic range metrics (geographic range size, occupied area, patch size distribution, geographic range filling, patch shape complexity, geographic range fractality) calculated from distribution data in the Atlas of Florae Europaeae rasterized using 10 x 10 km resolution maps. Dots represent individual species. Sample size was 827 species of plants, except the geographic range fractality where only 129 species with more than 9 patches were considered. Numbers indicate Pearson’s r correlation coefficient for linear, and Spearman’s Rho correlation coefficient for unimodal relationships. Note that only meaningful relationships should be interpreted.

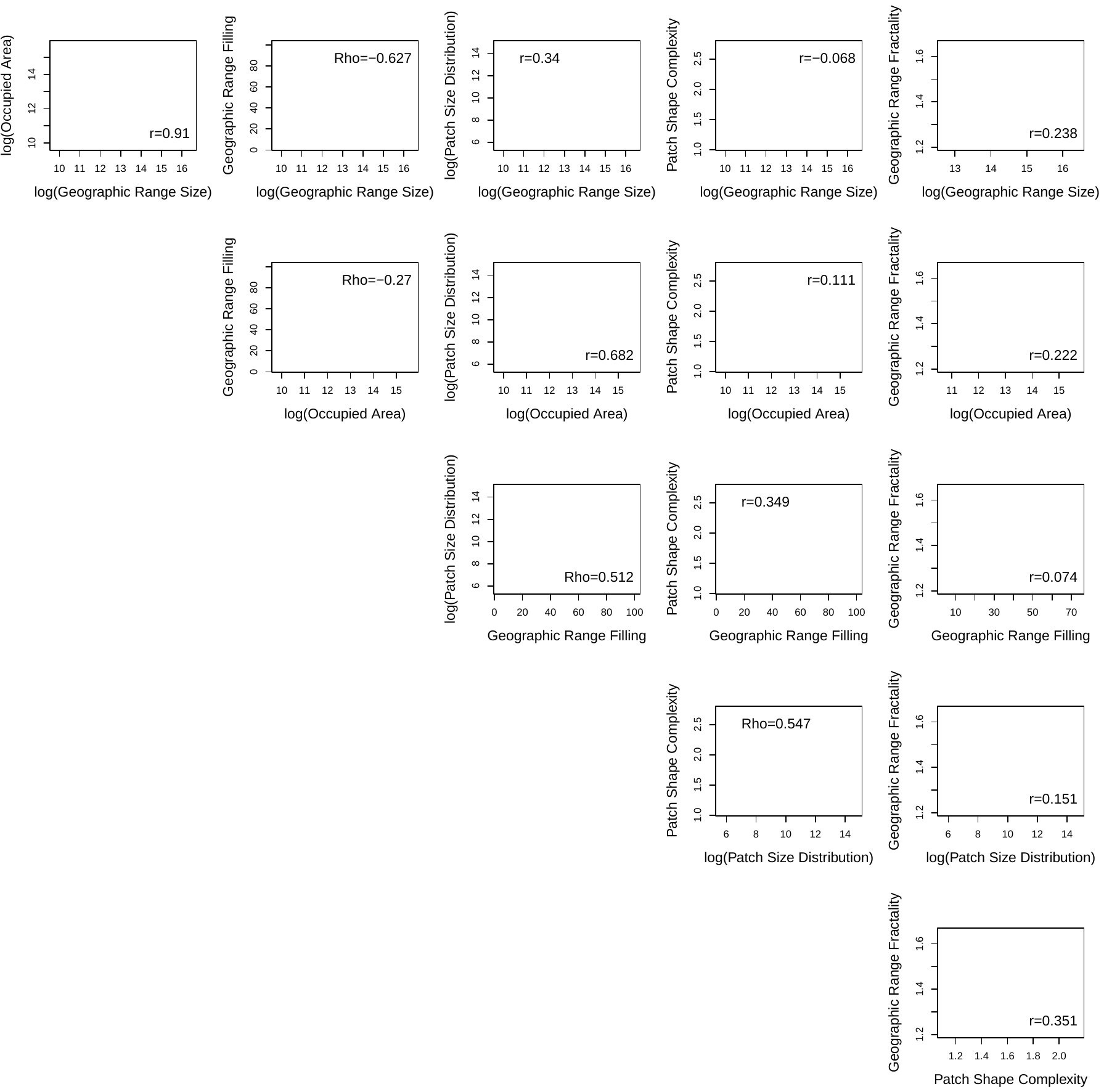

**S2.3 Sensitivity analyses**

**S2.3.1 Sensitivity analysis of the geographic range metrics to map resolution**

The geographic range size and structure metrics introduced here are insensitive to the grain (resolution) of the raster map i.e., on the number of grid cells within the habitat patches. In a theoretical example, the geographic range structure metrics were identical when calculated from raster maps at a resolution of 10 km and 1 km respectively (Figure S2.3.1.1).

cell size: 1 km

cell size: 10 km

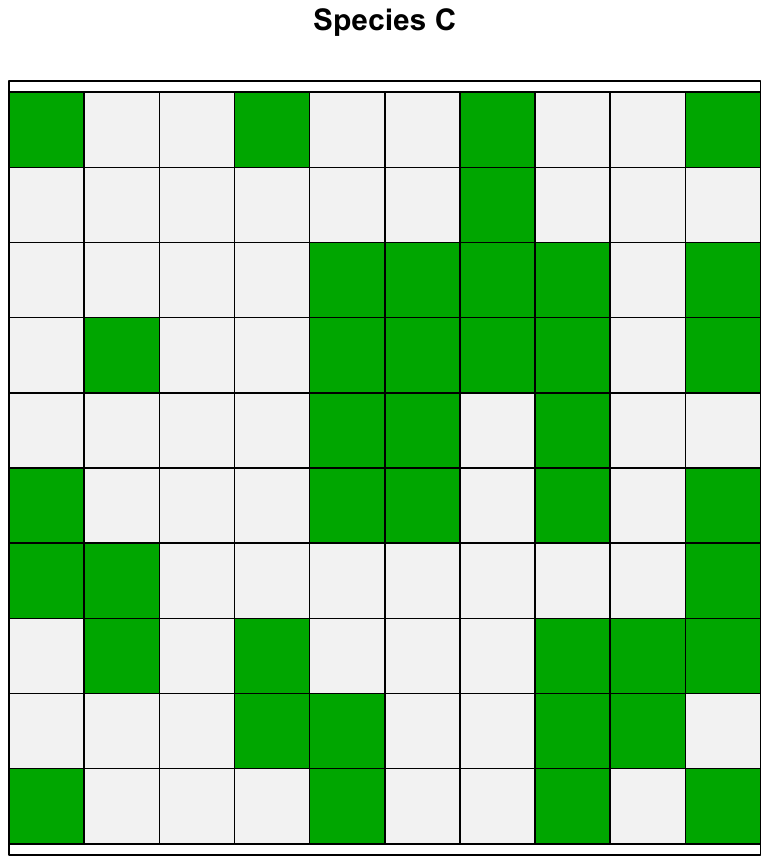

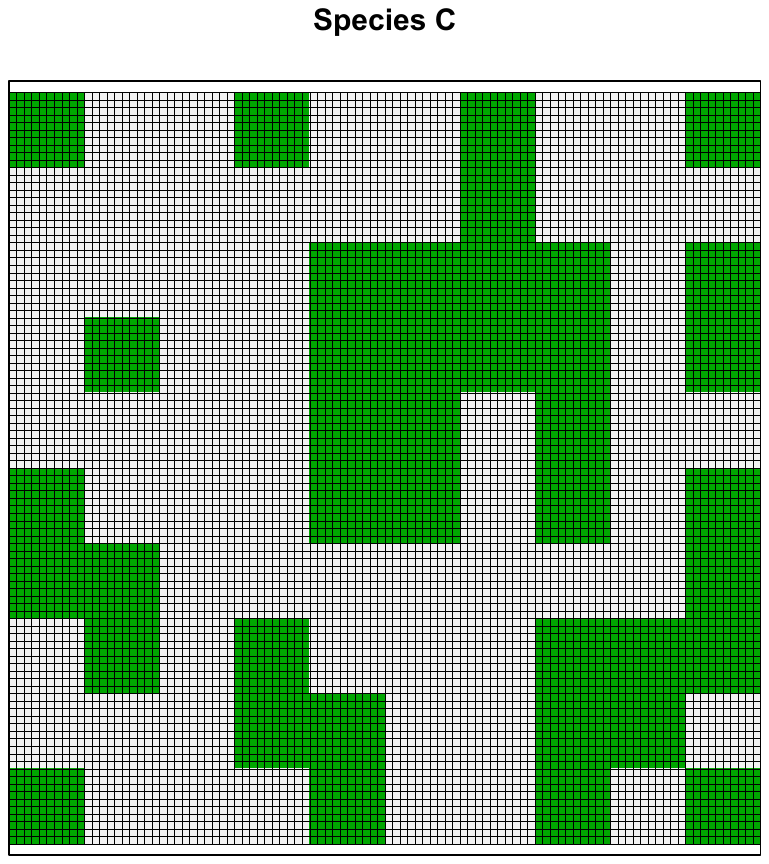

| **Geographic Range Structure metric** | **Resolution 10 km** | **Resolution 1 km** |
| --- | --- | --- |
| Geographic Range Size (km^2^) | 10000 | 10000 |
| Occupied Area (km^2^) | 4000 | 4000 |
| Patch Size Distribution (km^2^) | 378 | 378 |
| Geographic Range Filling (%) | 40 | 40 |
| Patch Shape Complexity (no unit) | 1.2 | 1.2 |
| Geographic Range Fractality (no unit) | 1.4 | 1.4 |

**Figure S2.3.1.1** The geographic range structure metrics calculated from identical maps at two different resolutions.

**S2.3.2 Sensitivity analysis of geographic range metrics to rasterization approach of the Atlas of Flora Europaea (AFE) grid**

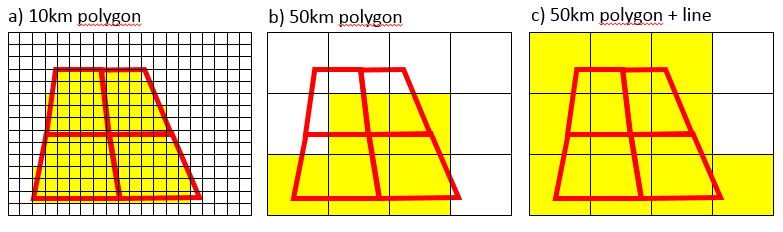
Rasterization of occurrence data available in different grid formats could affect the geographic range metrics. The occurrence data Atlas of Florae Europaeae has a resolution of approximately 50×50 km, and is available in a Common European Chorological Grid Reference System, CGRS)^[1]^. We rasterized the grid cells following three methods, each of which presented particular challenges: (**a**) using a raster map with a resolution lower than the width or length of the smallest AFE polygon cell (10 km) - the process is slow for large maps and a large number of species, and occasionally the grid cells were rasterized as “tetris” shapes; (**b**) using a raster with the width and length of the AFE polygon cells (50 km) – the process is faster but the cells that didn’t overlap with the raster cell center were not rasterized and we suspected that the ranges were underestimated; (**c**) using a 50 km polygon shapefile and a line shapefile, which uses 50 km cells but in addition this method rasterizes any pixel that the line touches, in this case we suspected that the resulting file overestimated the distribution (Figure S2.3.2.1). The geographic range metrics calculated with the 10 km polygon and 50 km polygon + line approaches were highly correlated (Pearson’s r > 0.9) for area metrics and range division metrics, and poorly correlated (Pearson’s r < 0.5) for patch shape metrics (Figure S2.3.2.2). However, the area and range division metrics were overestimated with the coarser method (most species fell above the red diagonal line. The patch shape metrics were also affected by the rasterization approach, with the coarse and less precise method producing lower patch shape complexity and lower range fractality than the finer 10 x 10 km resolution raster files. We therefore preferred to use the range structure metrics from maps rasterized with 10 x 10 km polygon shapefiles.

**Figure S2.3.2.1** Visual representation of the three different rasterization methods followed: (a) 10 km polygons (b) 50 km polygons and (c) 50 km polygons and a line shapefile.

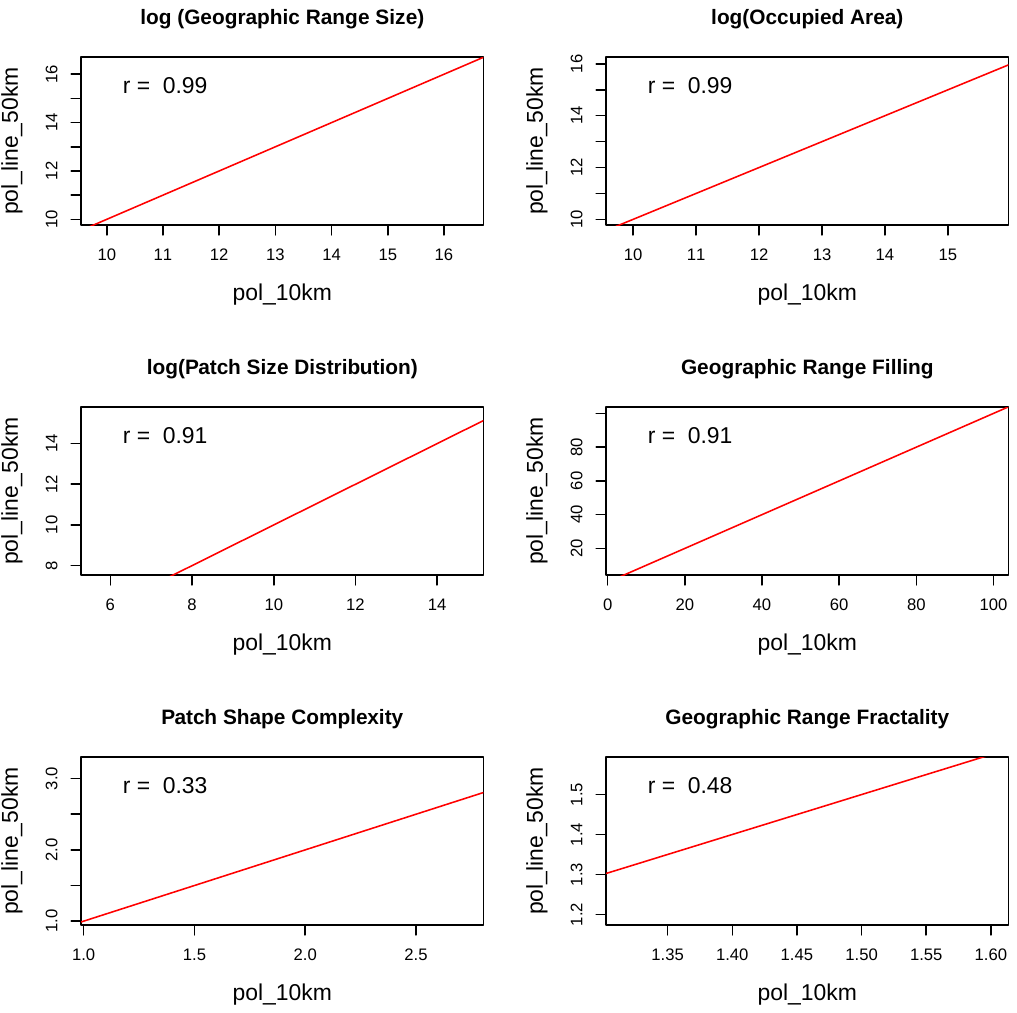

**Figure S2.3.2.2** Scatterplots showing correlations between six geographic range metrics calculated from maps rasterized using 10 km polygons (pol_10 km) and 50 km polygons + lines (pol_ine_50 km). Each dot represents a species. Sample size equals 784 species overlapping in the two datasets, except for geographic range fractality with 57 overlapping species with number of patches >9. Numbers represent the Pearson’s r correlation coefficient. The red diagonal represents the 1:1 values.

**S2.3.3 Sensitivity of geographic range metrics to approaches taken to estimate species’ geographic range size (GRS)**

We approximated species’ geographic range size (GRS) using a minimum convex polygon around 95% of occupied grid cells, to avoid overestimating the range size due to highly disjunct areas. We tested how this choice to define he minimum convex polygon may have influenced the geographic range structure metrics by comparing the geographic range structure metrics for geographic ranges defined using the minimum convex polygon around **(a)** 95% and **(b)** 100% of the occupied grid cells. We ran this test on occurrence data rasterized using the 50 km polygons + line approach, because at this resolution the choices affected more the area of the minimum convex polygon than when using fine (10km) resolution maps. The metrics calculated with the two approaches were highly correlated (Pearson’s r >0.9) and didn’t deviate much from the 1:1 diagonal line (Figure S2.3.3.1). Consequently, even at this resolution over- or underestimating a species’ geographic range size by omitting a few occurrence data doesn’t have a large effect on other geographic range metrics.

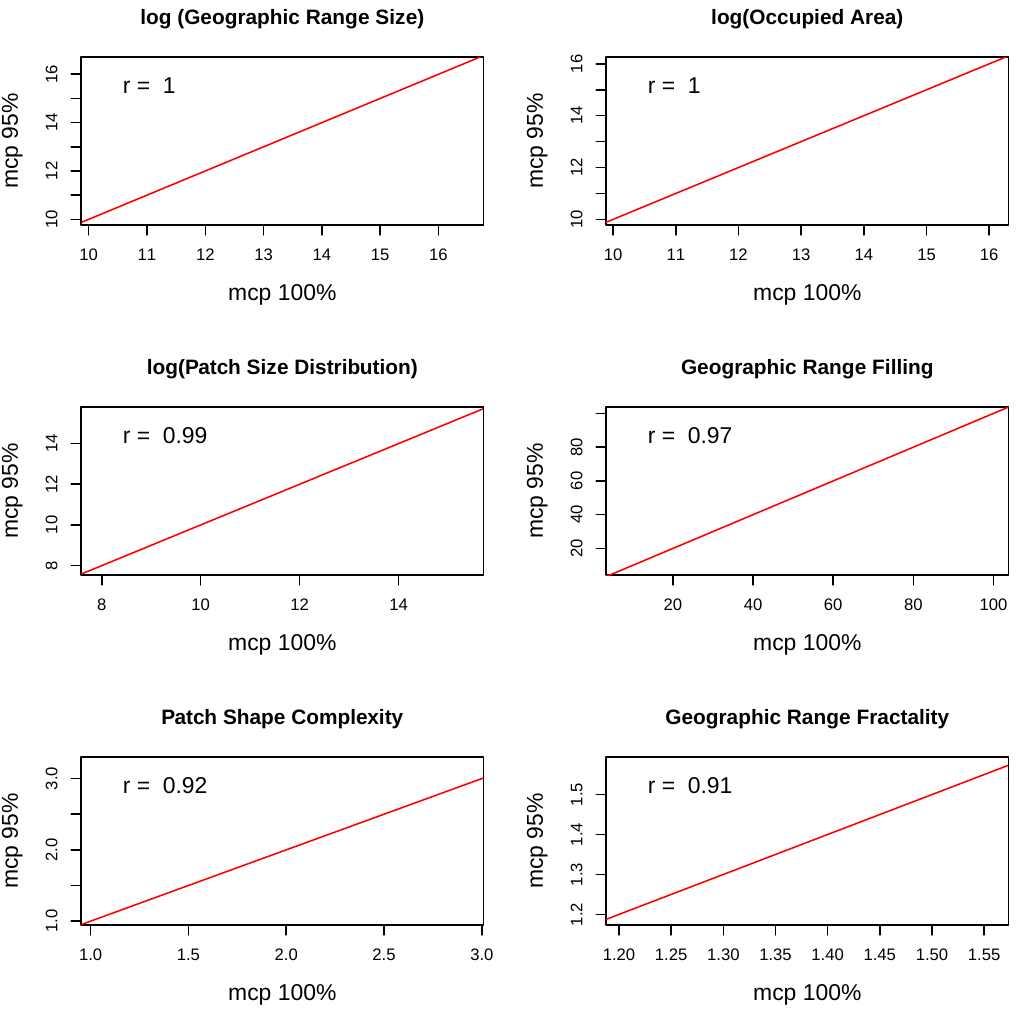

**Figure S2.3.3.1** Scatterplots showing correlations between six geographic range metrics calculated from geographic ranges defined using minimum convex polygons around 95% (axis y) and 100% (axis x) of the occupied grid cells. Dots represent individual species. Sample size equals 809 species that overlapped in the two datasets, except for the geographic range fractality with 64 overlapping species in the two datasets (species with number of patches > 9). Red diagonals represent 1:1 values. Numbers represent Pearson’s r correlation coefficient.

**S2.3.4 Sensitivity analysis of geographic range metrics to species distribution modeling technique**

To examine to what extent the geographic range metrics are influenced by the species distribution modeling approach taken, we compared the distribution patterns resulted from our main modeling approach (Ensemble of Small Models, ESM; Breiner et al. 2015) with those resulted from classical species distribution models based on ensemble technique (SDM; Thuiller et al. 2009; Appendix S1). We suspected that SDMs for a number of species could suffer from overfitting issues due to a number of predictors higher than 1/10 of the number of occurrences.

The metrics calculated from maps produced with the two modeling techniques were highly correlated for geographic range size, occupied area and patch size distribution (Pearson’s r ≥ 0.9), moderately strongly correlated for the geographic range filling (Pearson’s r > 0.7) and poorly correlated for the two patch shape metrics (Pearson’s r < 0.5) (Figure S2.3.4.1). For a large number of species, SDMs tended to underestimate species’ geographic range size, occupied area, patch size distribution and geographic range filling (many points fell under the 1:1 diagonal line).

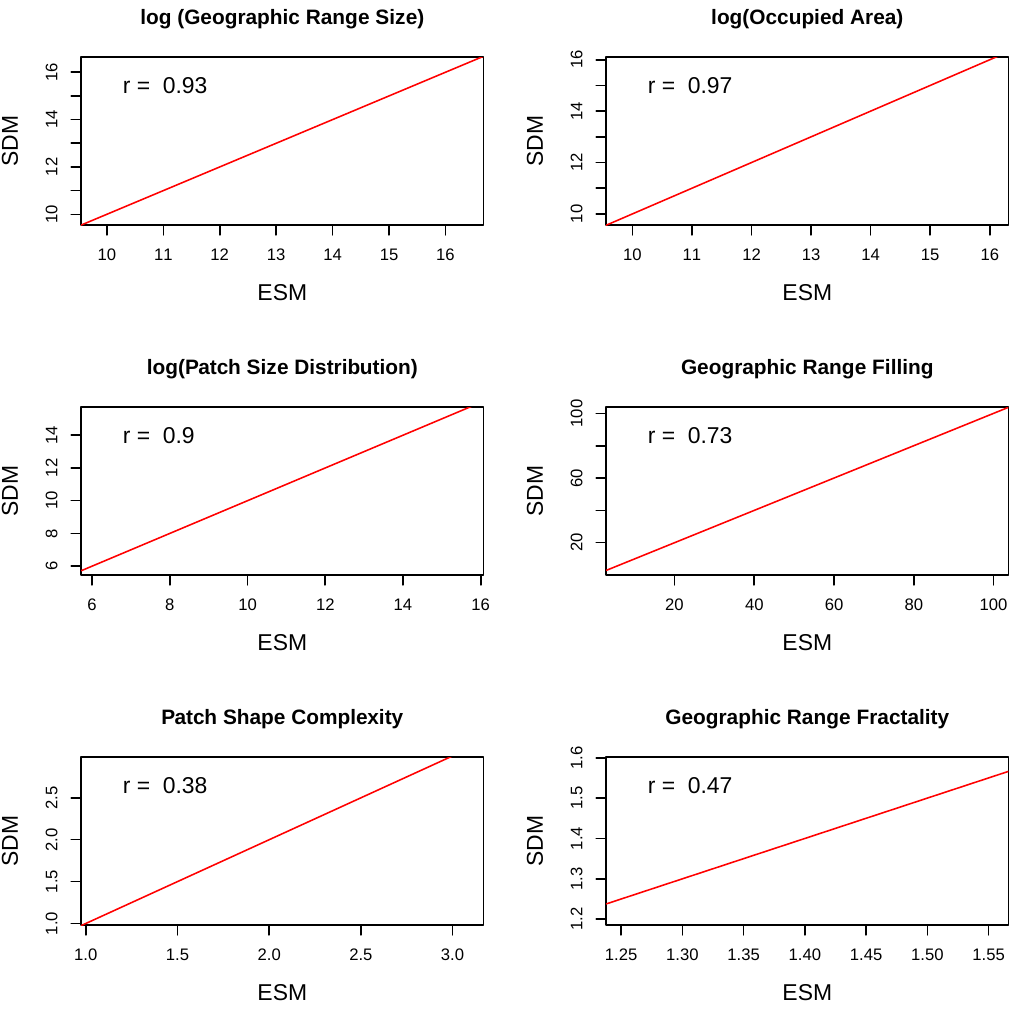

**Figure S2.3.4.1** Scatterplots showing correlations between six geographic range metrics calculated from maps based on two different modeling methods: Ensemble of Small Models (ESM) and classical species distribution models based on ensemble technique (SDM). Dots represent individual species. Sample size was 827 species for all metrics except geographic range fractality with 177 species. For this analysis we used maps rasterized at 10 x 10 km resolution.

**S2.3.5 Sensitivity analysis of geographic range metrics to prediction error of species distribution models**

One of the main sources of uncertainty in predictive modeling of species distribution stems from the method used to transform probabilities of occurrence produced by models into binary predictions of species presence and absence (Nenzen & Araujo 2011). To test the sensitivity of geographic range metrics to prediction error, we binarized the continuous predictions using thresholds in habitat suitability corresponding to maxKappa, maxTSS, sensitivity=specificity, and MinROCdist as in Nenzen & Araujo (2011), and we compared the geographic range metrics calculated from the four maps for each species (Figure S2.3.5.1). We used habitat suitability maps predicted by the Ensemble of Small Models approach (Breiner et al. 2015) at a resolution of 10 x 10 km for over 800 species in Europe. All geographic range metrics using MaxTSS were the least correlated with the metrics calculated using MaxKappa, and they were highly correlated with range metrics calculated using Se=Sp and MinROCdist (Figure S2.3.5.2).

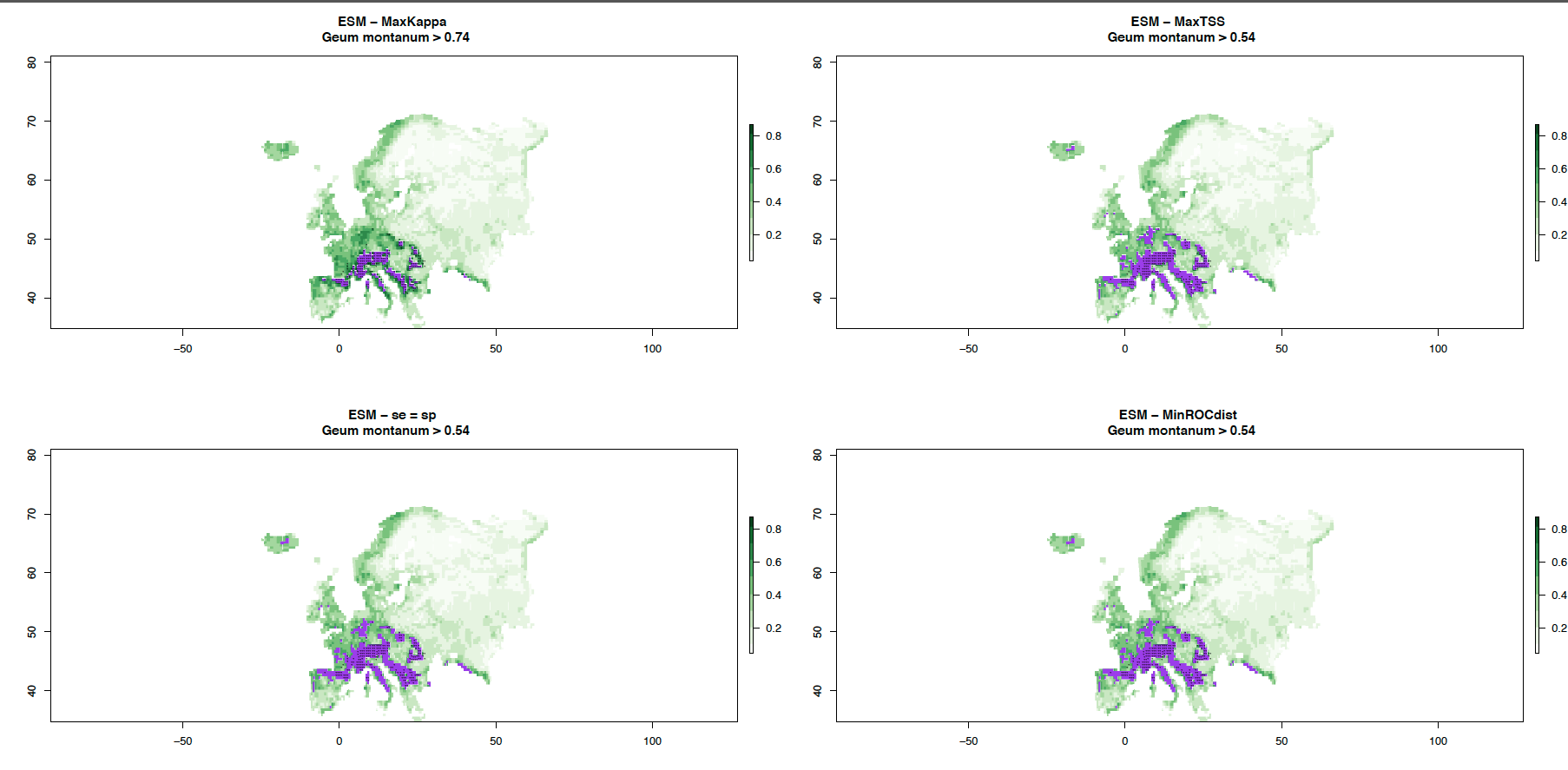

Figure **S2.3.5.1** Predicted distributions for an example species (*Geum montanum*) using Max Kappa, maxTSS, Sensitivity = Specificity, and MinROC distance.

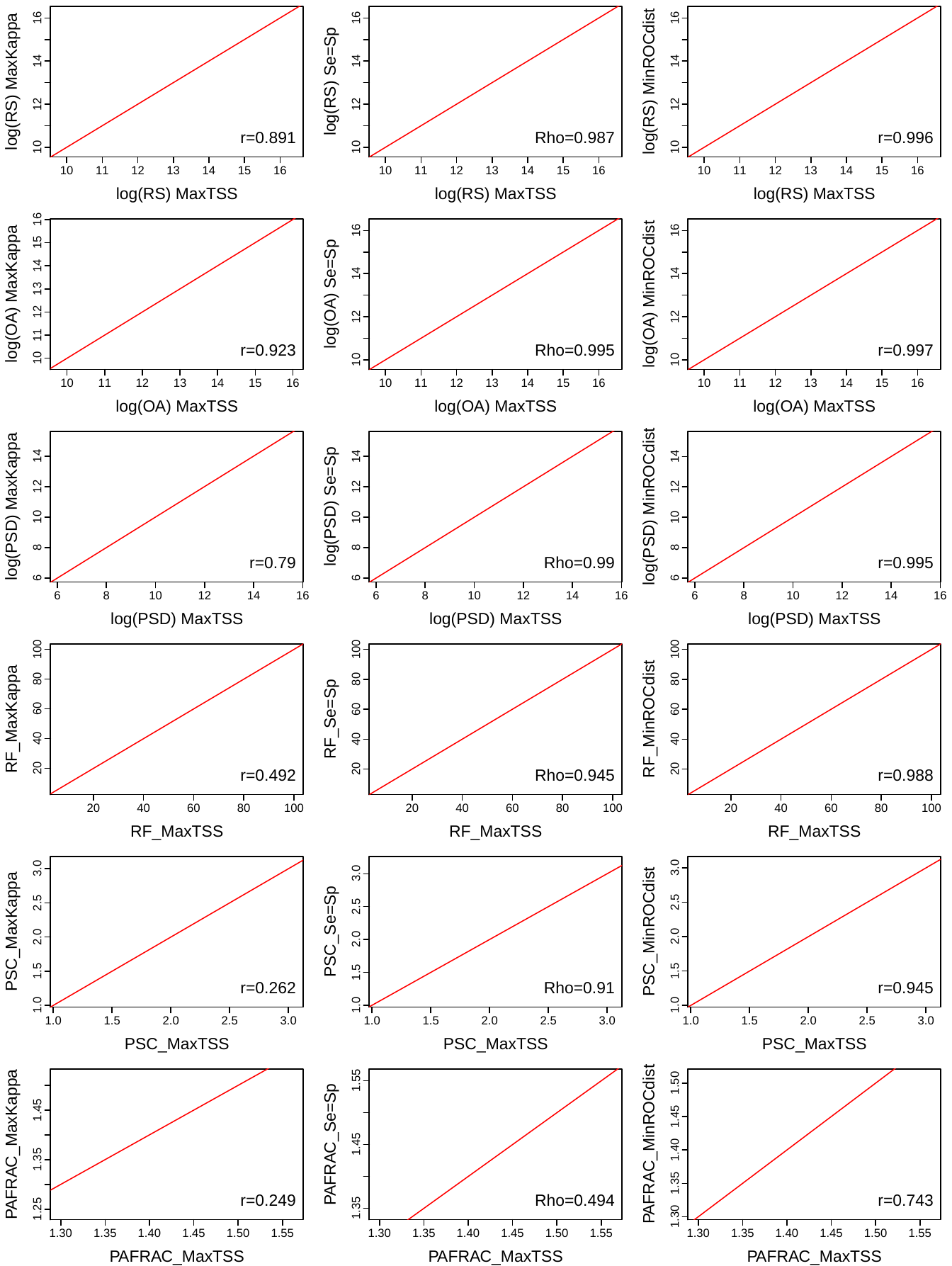

**Figure S2.3.5.2** Scatterplots showing correlations between geographic range metrics calculated from maps based on predictions using maxTSS and maps based on predictions using maxKappa, Sensitivity=Specificity, and MinROCdist. Dots represent individual species. RS = geographic range size, OA = occupied area, PSD = patch size distribution, RF = geographic range filling, PSC = patch shape complexity, PAFRAC = geographic range fractality. Sample size equals 827 species for all metrics except geographic range fractality with 41 species for which the number of habitat patches was higher 9 across all maps.

**S2.3.6 Sensitivity of geographic range metrics to habitat quality thresholds**

We examined how geographic range patterns are affected by increasing the thresholds used to define habitat quality. We used habitat suitability maps predicted by the Ensemble of Small Models approach (Breiner et al. 2015) at a resolution of 10 x 10 km. We used six thresholds to draw six binary maps with increasing predicted habitat quality corresponding to the 0, 0.1, 0.2, 0.3, 0.4 and 0.5 quantiles of the habitat suitability values predicted for all occupied grid cells within a species’ range (modified from Engler et al. 2004) (Figure S2.3.6.1). The map with the largest surface encompassed all habitat suitability values predicted for occupied grid cells i.e., it included a large range of habitat suitability values, while the map with the smallest surface encompassed the best 50% of the habitat suitability values predicted for occupied grid cells.

To explore whether changes in geographic range size and structure with increasing habitat suitability threshold were significant, we fitted linear mixed models with the metric of interest as predicted variable, and habitat suitability threshold as explanatory variable. Habitat suitability threshold was introduced as fixed effect and species as random effect. Habitat patches of increasing habitat suitability were smaller in size, less continuous, they occupied a smaller proportion of the species’ range, they were more irregular, and larger patches were much more complex than smaller patches (Figure S2.3.6.2).

**Figure S2.3.6.1** Predicted area for *Geum montanum* in Europe, for six different habitat suitability thresholds. The green shades indicate habitat suitability as predicted by the Ensemble of Small Models for both occupied and non-occupied grid cells. Purple color indicates grid cells above the specific quantile thresholds: Q0=0, Q10=0.1, Q20=0.2, Q30=0.3, Q40=0.4 and Q50=0.5. The area of the purple maps decreases according to the increasing habitat suitability threshold identified by the six habitat suitability quantiles. Note that both occupied and non-occupied grid cells can be included as suitable.

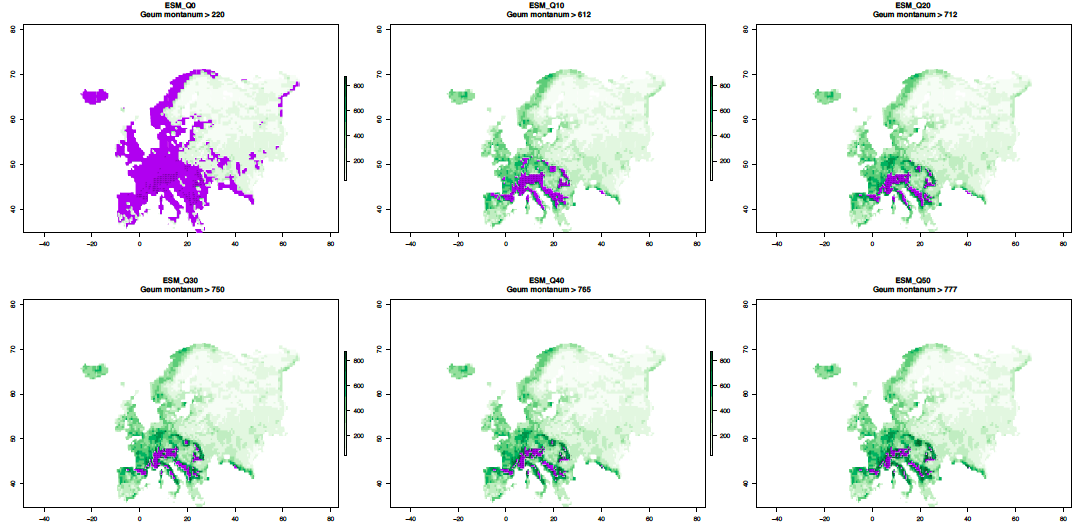

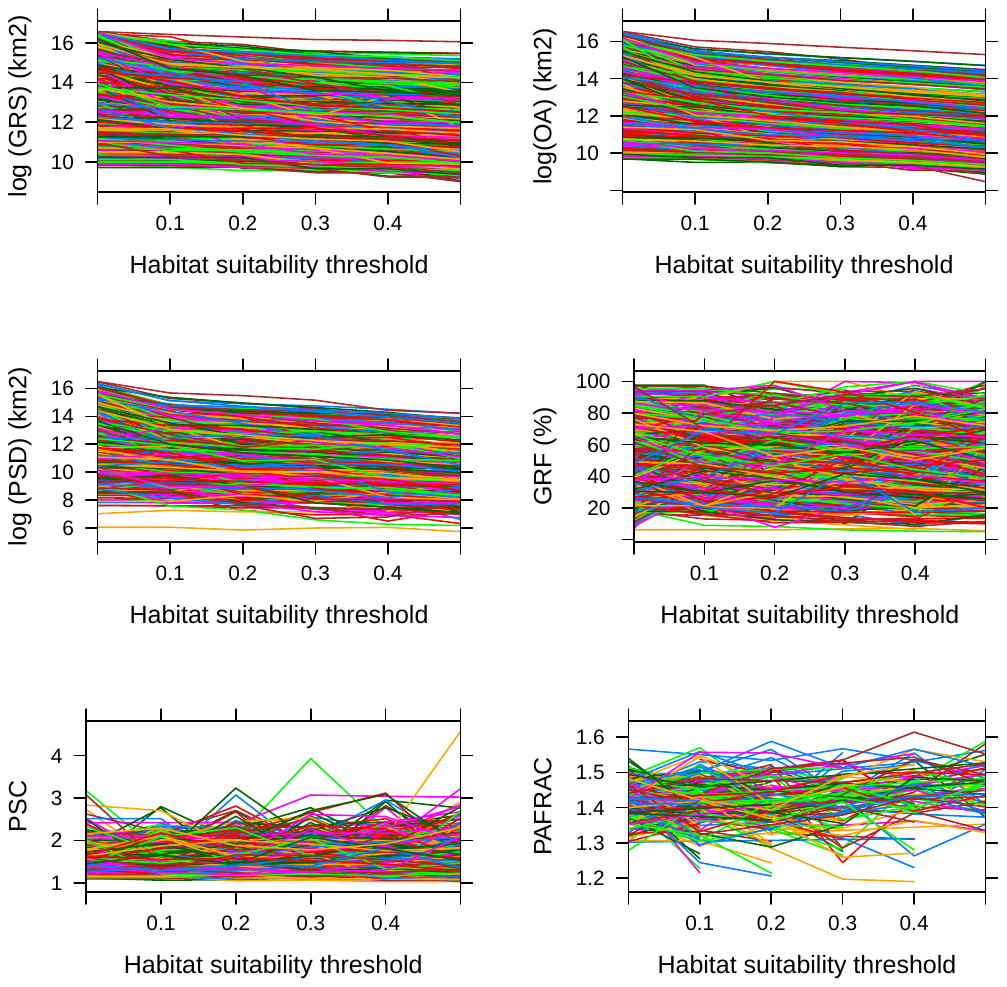

**Figure S2.3.6.2** Relationship between six geographic range metrics and habitat suitability threshold for 811 species endemic to Europe (except PAFRAC with 332 species). The parameters estimated by linear mixed model were: GRS (geographic range size, β = -3.027±0.035 SE, p<0.001), OA (occupied area, β = -3.066±0.027 SE, p<0.001), PSD (patch size distribution, β = -3.173±0.045 SE p<0.001), GRF (geographic range filling, β = -2.161±0.952 SE, p=0.023), PSC (patch shape complexity, β = 0.042±0.021 SE, p=0.051), PAFRAC (geographic range fractality, β = 0.039±0.014 SE, p=0.005).

**S3. Details of PCA analysis**

The PCA analysis was based on the same eigth climate variables used in the species distribution modeling: *tdr.max* (mean diurnal temperature range, Bio02, maximum), *tar.min* (temperature annual range, Bio07, minimum), *twetq.min* (mean temperature of wettest quarter, Bio08, minimum), *tdryq.max* (mean temperature of driest quarter, Bio09, maximum), *twarmq.max* (mean temperature of warmest quarter, Bio10, maximum), *p.mean* (annual precipitation, Bio 12, mean), *ps.max* (precipitation seasonality, Bio15, maximum), and *pdryq.min* (precipitation of driest quarter, Bio17, minimum). The first two PCA axes explained 54.0% of variance in the environmental space of the species in our dataset (Figure S3.1).

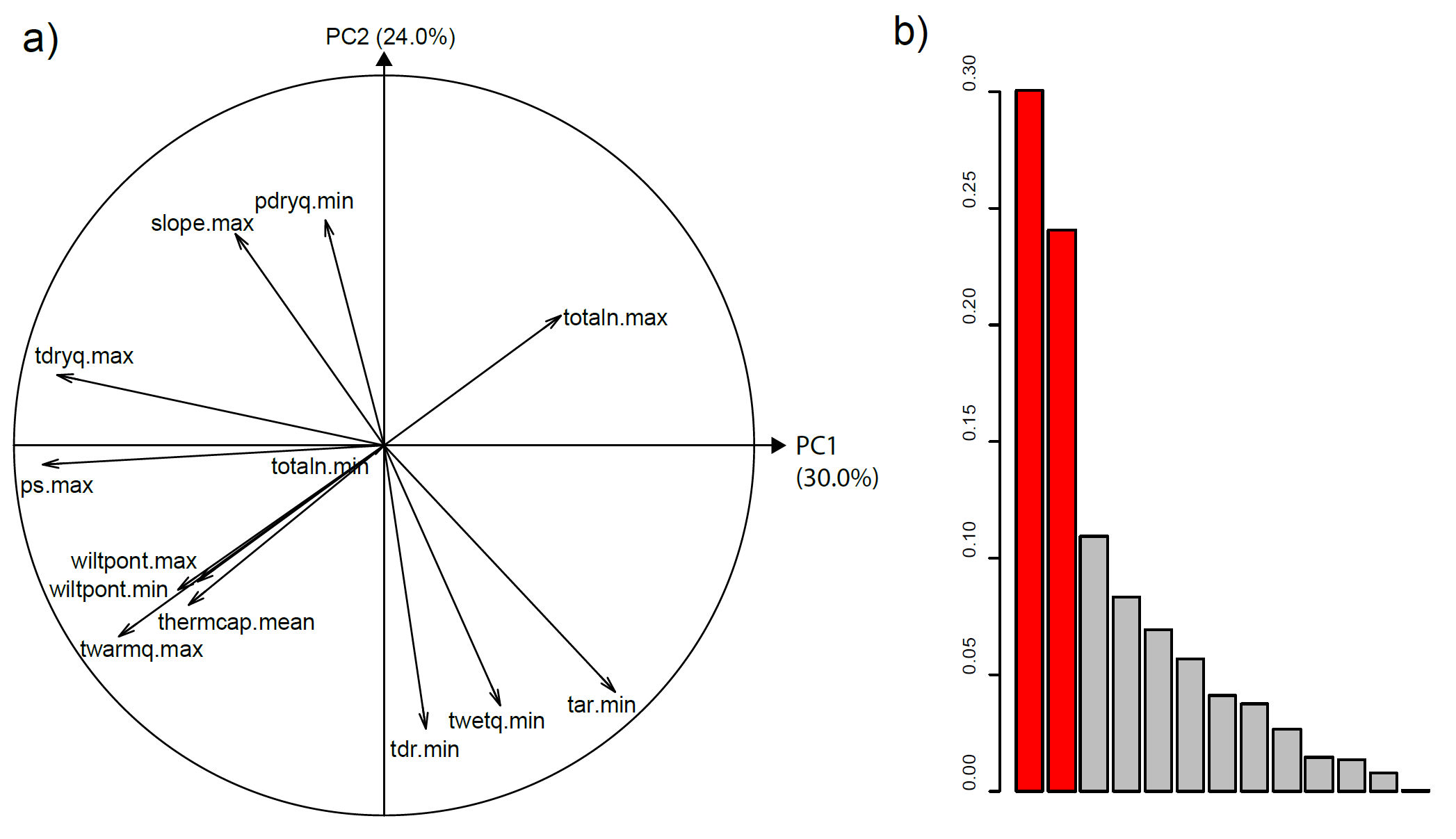

**Figure S3.1.** Results of the PCA analysis showing **a)** the loadings of each climate variable on the first two PCA axes and **b)** the percent variance explained by each PCA axes, with the first two axes highlighted in red.

**S4. Results of statistical and phylogenetic analyses of geographic range metrics.**

**S4.1 Comparisons of geographic range metrics quantified from observed and predicted distribution maps**

**Table 4.1.1** Results of the paired *t*-tests and phylogenetically corrected paired *t*-tests between observed and predicted geographic range metrics. *p* values of significant differences are bolded.

|  | **Paired t-test** | |  | **Phylogenetically corrected t-test** | | | |
| --- | --- | --- | --- | --- | --- | --- | --- |
| **Range metric** | **t** | **df** | **p** | **t** | **df** | **p** | **lambda** |
| Geographic Range Size | -20.810 | 812 | **0.000** | -1.753 | 810 | 0.080 | 0.368 |
| Occupied Area\|Suitable Area | -19.116 | 812 | **0.000** | -1.429 | 810 | 0.153 | 0.734 |
| Patch Size Distribution | -11.459 | 812 | **0.000** | -0.498 | 810 | 0.619 | 0.803 |
| Geographic Range Filling | -5.897 | 812 | **0.000** | -5.931 | 810 | **0.000** | 0.000 |
| Patch Shape Complexity | -1.772 | 812 | 0.077 | -1.772 | 810 | 0.077 | 0.000 |
| Geographic Range Fractality | 0.555 | 70 | 0.581 | 0.564 | 68 | 0.575 | 0.000 |

**S4.2 Phylogenetic signal in species’ geographic range size and structure**

We constructed a phylogenetic tree for our species using the tree available in Zanne et al. (2014). We estimated the  phylogenetic relationships between species for each geographic range metric for occurrence data and predicted habitat maps using Pagel’s λ (1999) and Blomberg’s K (Blomberg et al. 2003). λ measured the similarity of covariances among species to covariances expected under Brownian motion, and K was interpreted as a measure of the partitioning of variance among and between clades.

**Table S4.2.1** Results of the tests of phylogenetic signal for the geographic range metrics calculated from observed and predicted distribution maps and the proportion between the predicted and observed range structure metric. Occurrence = observed distribution in the Atlas of Florae Europaeae, ESM = predicted distribution Ensemble of Small Models approach, Proportion = ESM/Occurrence. The phylogenetic signal was calculated using the *phylosig* command in ‘phytools’ package in R using two metrics: λ = Pagel’s λ, and K. p(λ) = significance test of λ, p(K) = significance test of K calculated using 1000 simulations. *p* values indicating significant phylogenetic relationships are bolded

| **Geographic Range metric** | **Data** | **Lambda** | **P(Lambda)** | **K** | **P(K)** |
| --- | --- | --- | --- | --- | --- |
| Geographic Range Size | Occurrence | **0.794** | **0.000** | 0.042 | 0.520 |
| Geographic Range Size | ESM | **0.731** | **0.000** | 0.050 | 0.156 |
| Occupied Area | Occurrence | **0.852** | **0.000** | 0.032 | 0.719 |
| Suitable Area | ESM | **0.804** | **0.000** | 0.044 | 0.404 |
| Patch Size Distribution | Occurrence | **0.744** | **0.000** | 0.016 | 0.910 |
| Patch Size Distribution | ESM | **0.807** | **0.000** | 0.033 | 0.677 |
| Geographic Range Filling | Occurrence | 0.151 | 1.000 | 0.042 | 0.260 |
| Geographic Range Filling | ESM | 0.000 | 1.000 | 0.039 | 0.350 |
| Patch Shape Complexity | Occurrence | 0.000 | 1.000 | 0.031 | 0.759 |
| Patch Shape Complexity | ESM | 0.000 | 1.000 | 0.037 | 0.564 |
| Geographic Range Fractality | Occurrence | 0.000 | 1.000 | 0.074 | 0.924 |
| Geographic Range Fractality | ESM | 0.000 | 1.000 | 0.169 | 0.112 |

**S.4.3** pGLS model results showing the relationship between observed geographic range metrics and the log response ratio between metrics calculated from predicted and observed distributions. The first column shows the model structure with metric names as used in Fragstats, and full names of predicted variables. The next columns show the estimates, standard errors, and *p* values of the variables, the adjusted R squared value of the model and the maximum likelihood estimates of lambda and the corresponding *p* values of upper and lower bounds of the lambda.

| **Model structure and predicted variable** | **Variable** | **β** | **β(SE)** | **p** | **R2** | **lambda** | **p (lambda)** |
| --- | --- | --- | --- | --- | --- | --- | --- |
| *log(propRS) ~ log(range.size) + I(log(range.size)^2)* | Intercept | -28.656 | 1.651 | <0.001 | 0.289 | 0 | 1/<0.001 |
| Predicted/Observed Geographic Range Size | log(Range Size) | 4.564 | 0.260 | <0.001 |  |  |  |
|  | log(Range Size)^2 | -0.175 | 0.010 | <0.001 |  |  |  |
| *log(propTA) ~ log(total.area) + I(log(total.area)^2)* | Intercept | -31.239 | 1.334 | <0.001 | 0.480 | 0 | 1/<0.001 |
| Predicted/Occupied Area | log(Occupied Area) | 5.175 | 0.222 | <0.001 |  |  |  |
|  | log(Occupied Area)^2 | -0.205 | 0.009 | <0.001 |  |  |  |
| *log(propMESH) ~ log(effective.mesh.size)* | Intercept | 4.937 | 0.626 | <0.001 | 0.134 | 0.386 | 0.0006/<0.001 |
| Predicted/Observed Patch Size Distribution | log(Patch Size Distribution) | -0.355 | 0.032 | <0.001 |  |  |  |
| *log(propRF) ~ prop.landscape* | Intercept | 1.500 | 0.068 | <0.001 | 0.411 | 0 | 1/<0.001 |
| Predicted/Observed Geographic Range Filling | Geographic Range Filling | -0.050 | 0.003 | <0.001 |  |  |  |
|  | Geographic Range Filling^2 | 0.000 | 0.000 | <0.001 |  |  |  |
| *log(propSHAPE) ~ mean.shape.index* | Intercept | 0.747 | 0.030 | <0.001 | 0.427 | 0 | 1/<0.001 |
| Predicted/Observed Patch Shape Complexity | Patch Shape Complexity | -0.490 | 0.020 | <0.001 |  |  |  |
| *log(propPAFD) ~ perimeter.area.frac.dim* | Intercept | 1.198 | 0.084 | <0.001 | 0.746 | 0 | 1/<0.001 |
| Predicted/Observed Geographic Range Fractality | Geographic Range Fractality | -0.844 | 0.059 | <0.001 |  |  |  |

**S4.4** The effect sizes (slope and 95% confidence intervals) of niche breadth (Niche_br), median range latitude (Lat), median range longitude (Lon), topographic heterogeneity (SD elevation), and mean distance from coastline (Dist_coast) on six geographic range metrics: geographic range size (GRS), occupied (suitable) area (OA, SA), patch size distribution (PSD), geographic range filling (GRF), patch shape complexity (PSC), geographic range fractality (GRFRAC) modeled using pGLS on species’ observed and predicted distributions. Positive slope values indicate an increase, and negative slope values indicate a decrease of each response variable with increasing values of the explanatory variables. Confidence intervals that encompass 0 indicate non-significant effects. Model details are presented in S4.5 below.

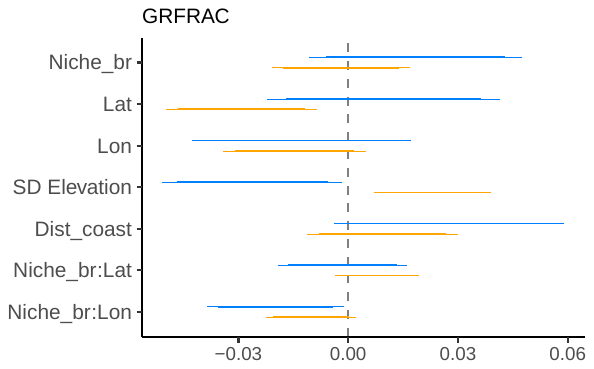

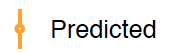

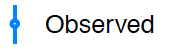

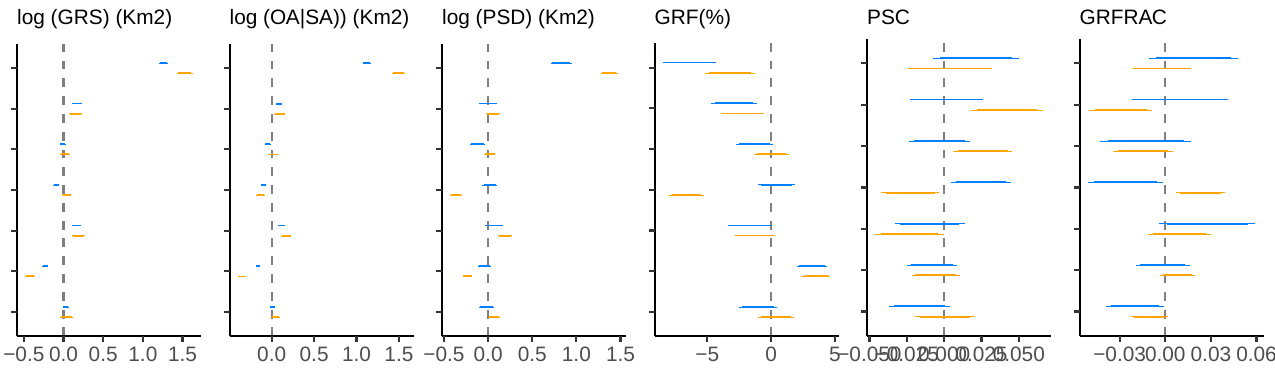

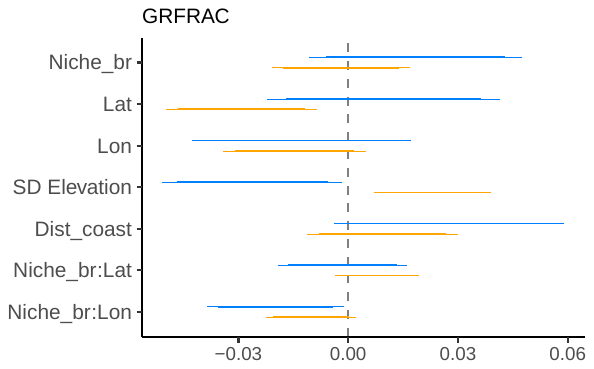

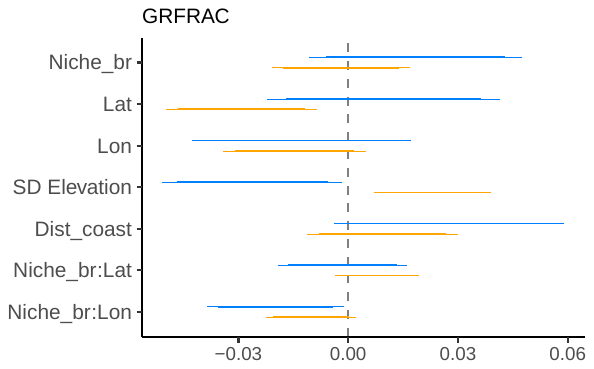

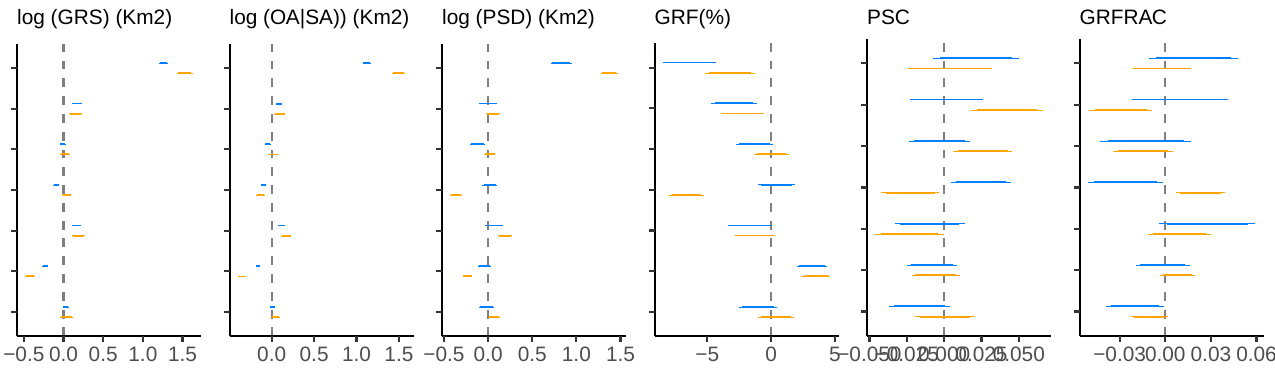

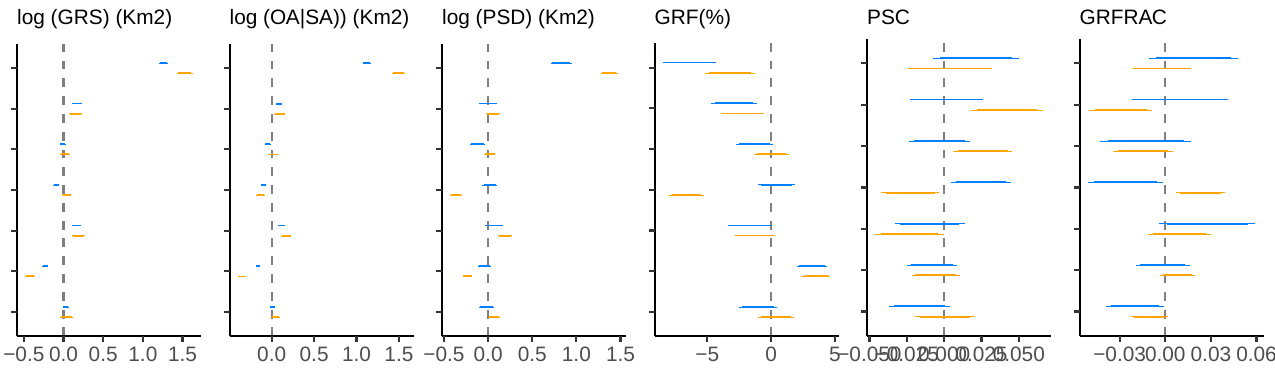

**S4.5** Scatterplots showing the effect of topographic heterogeneity (SD elevation), niche breadth, median range latitude, median range longitude and mean distance from continental coastline on six geographic range metrics: geographic range size, occupied (suitable) area, patch size distribution, geographic range filling, patch shape complexity, geographic range fractality modeled using pGLS on species’ observed and predicted distributions. Model fit plots are shown on the right. The first column of the table shows the full names of predicted variables. The next columns show estimates, standard errors, and *p* values of variables, adjusted R squared value of the model and maximum likelihood estimates of lambda and the corresponding *p* values of upper and lower bounds of lambda.

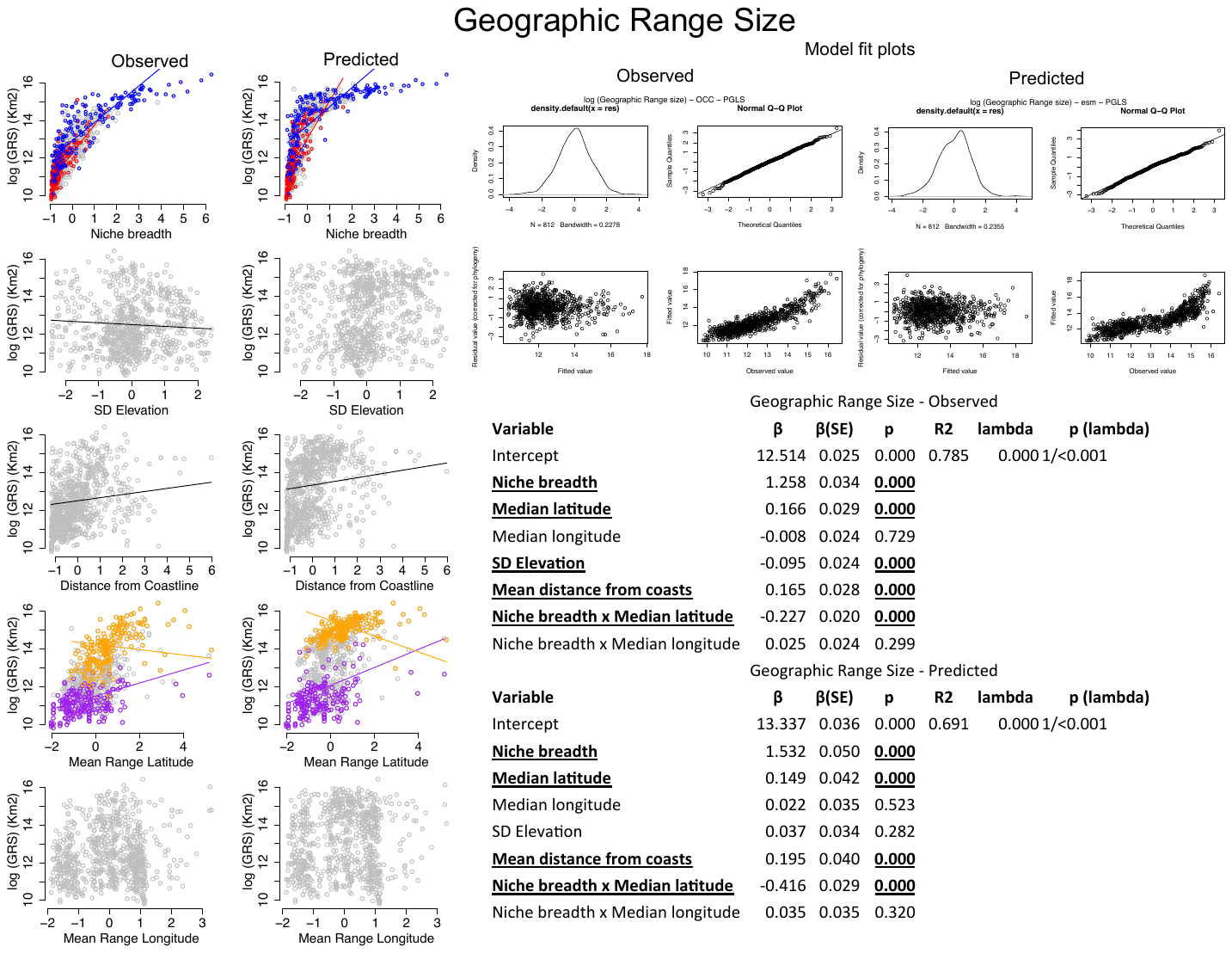

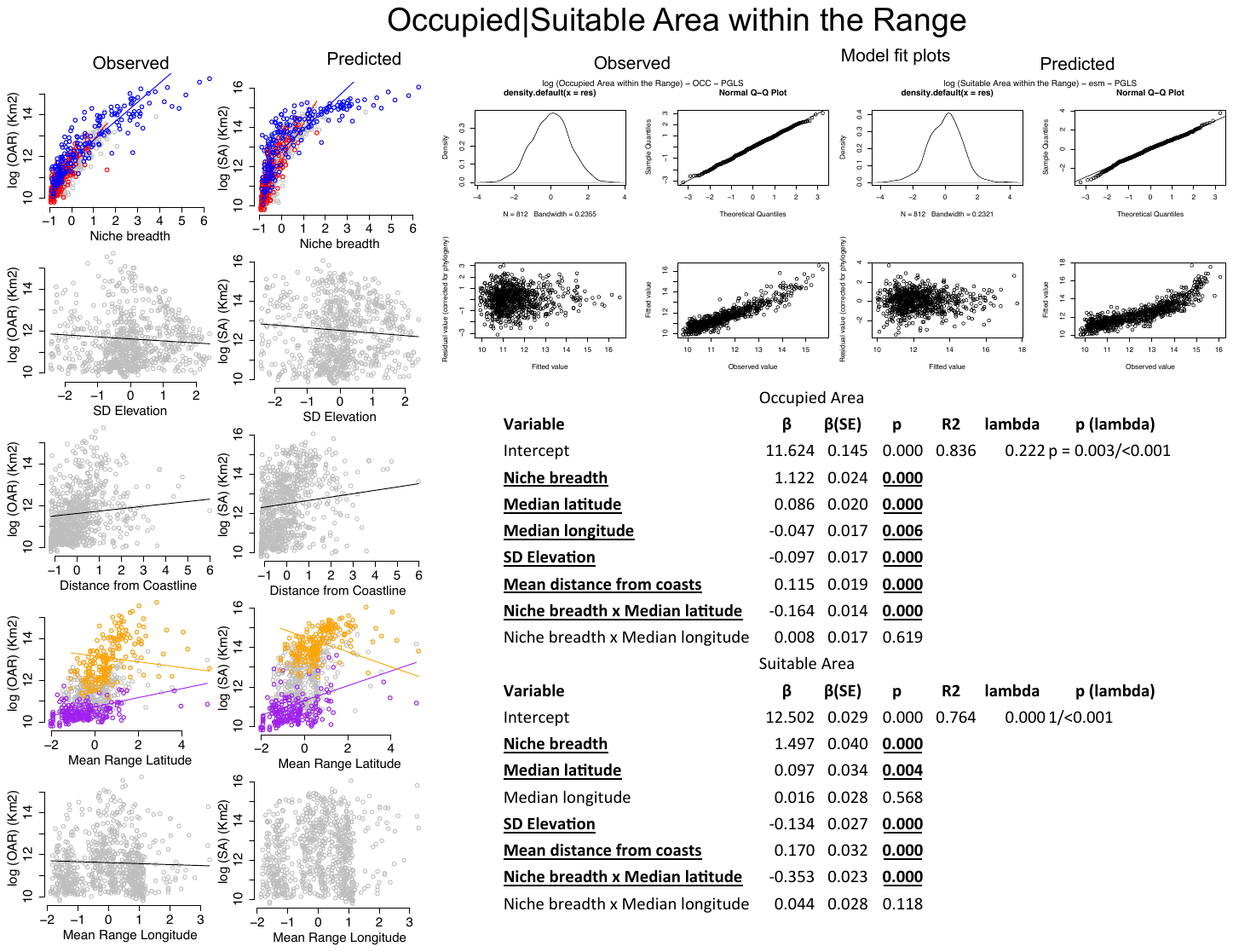

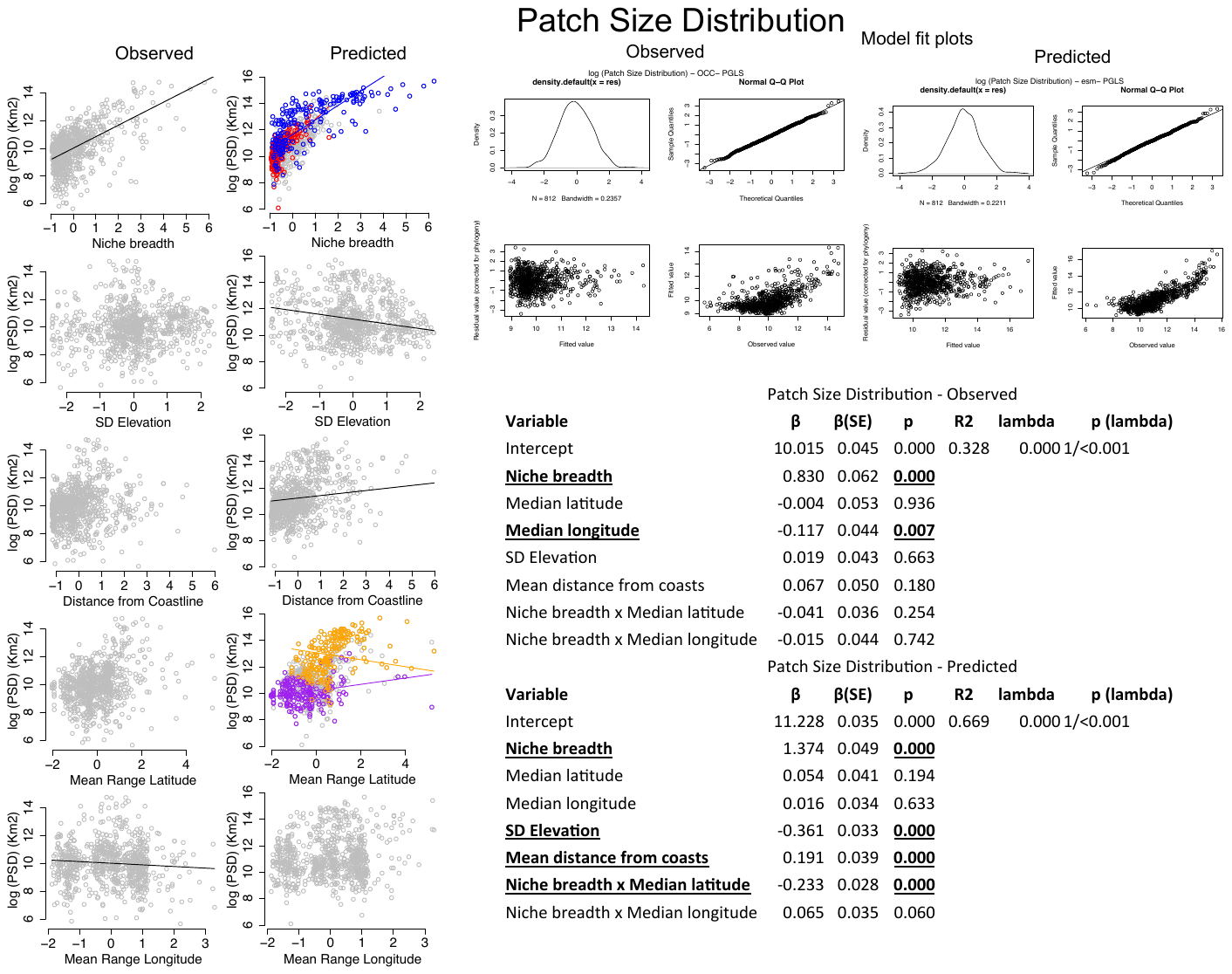

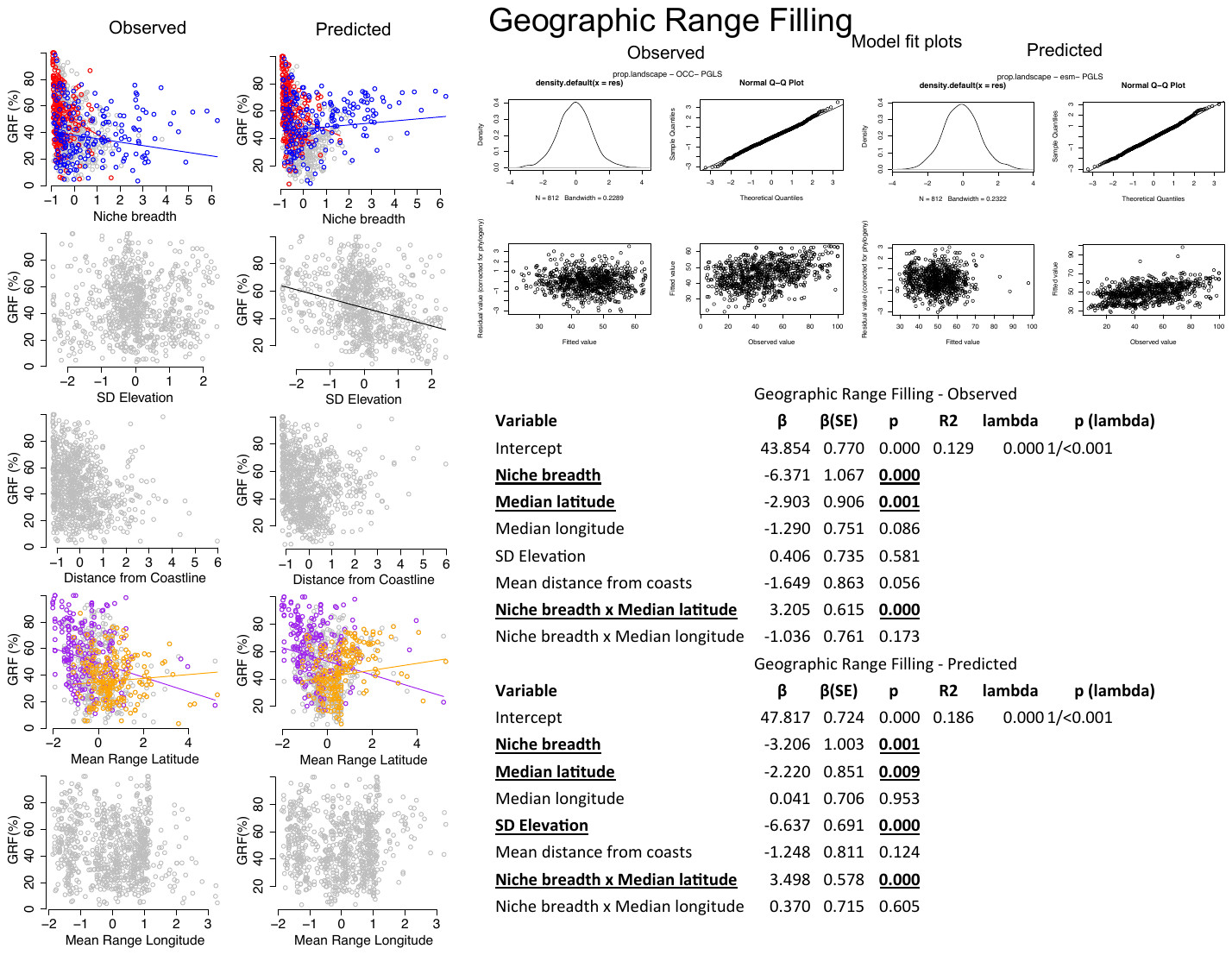

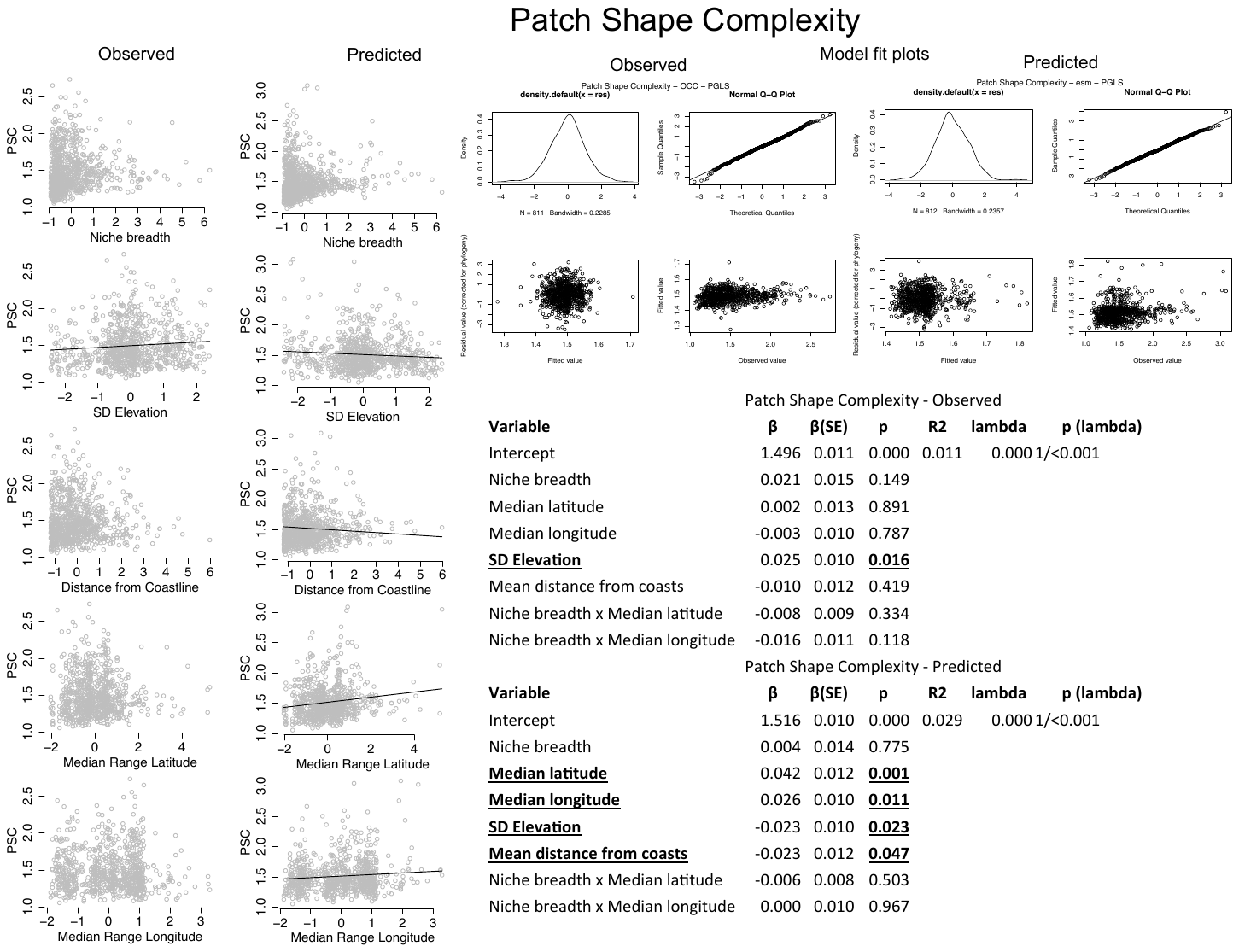

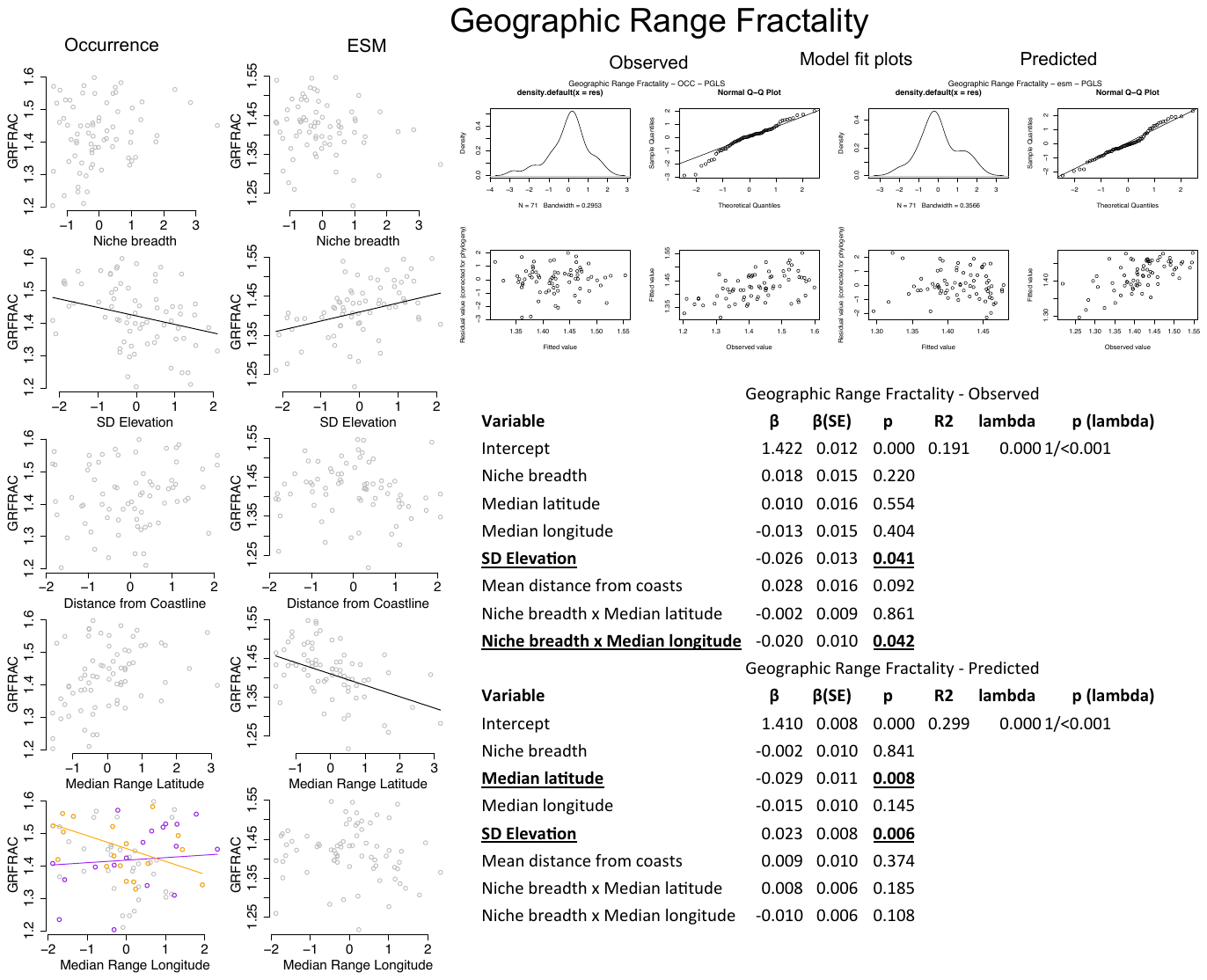

1. https://www.eea.europa.eu/data-and-maps/data/common-european-chorological-grid-reference-system-cgrs
